# Fission yeast polycystin Pkd2p promotes the cell expansion and antagonizes the Hippo pathway SIN

**DOI:** 10.1101/2021.05.21.444707

**Authors:** Debatrayee Sinha, Denisa Ivan, Ellie Gibbs, Madhurya Chetluru, John Goss, Qian Chen

## Abstract

Polycystins are conserved mechanosensitive channels whose mutations lead to the common human renal disorder ADPKD. Previously we discovered that the plasma membrane-localized fission yeast homologue Pkd2p is an essential protein required for cytokinesis, but the mechanism remains unclear. Here, we isolated a novel temperature-sensitive mutant *pkd2-B42.* Among its strong growth defects, the most unique was that many mutant cells often lost significant portion of their volume in just 5 minutes followed by a gradual recovery, a process that we termed Deflation. Unlike cell lysis, deflation did not result in the plasma membrane rupture and it occurred independently from the cell cycle progression. The tip extension of *pkd2-B42* cells was 80% slower than the wild type and their turgor pressure was 50% lower. Both *pkd2-B42* and the other mutant *pkd2-81KD* partially rescued the mutants of the yeast Hippo signaling pathway Septation Initiation Network, by preventing cell lysis, enhancing septum formation, and doubling the number of Sid2/Mob1 molecules at the spindle pole bodies. We conclude that Pkd2p promotes cell size expansion during interphase by regulating turgor pressure and antagonizes SIN during cytokinesis.

**Summary statement:** Mutations of polycystins lead to human genetic disorder ADPKD. We discovered that the fission yeast homologue Pkd2p promotes the cell expansion during interphase growth and antagonizes the Hippo pathway SIN during cytokinesis.

## Introduction

Cytokinesis is the last stage of cell division when two daughter cells separate. It initiates the assembly of an actomyosin contractile ring at the medial division plane. The ring provides the mechanical force required for the cleavage furrow ingression, eventually leading to cell separation. Fission yeast, *Schizosaccharomyces pombe,* represents an excellent model organism to study this evolutionally conserved cellular process. A large number of cytokinetic genes have been identified in this unicellular eukaryote through genetic screens. Cell biology studies have helped to reveal their functions, most of which are likely conserved in metazoans (for review, see (Pollard and Wu, 2010)).

In fission yeast cytokinesis, the Hippo signaling pathway SIN (septation initiation network) plays an essential role. This differs slightly from the metazoan Hippo pathway which modulates cell growth and organ size (Camargo et al., 2007; Dong et al., 2007). Nevertheless, both consist of a highly conserved kinase cascade (for reviews, see (Johnson et al., 2012; Simanis, 2015; Yu et al., 2015)). At the top of the fission yeast SIN pathway is a GTPase Spg1p, localized at the spindle pole bodies (SPBs) throughout mitosis (Schmidt et al., 1997). Its activation propagates the SIN signal through a chain of three kinases including the essential nuclear Dbf2-related (NDR) family kinase Sid2p, the homologue of human LATS kinase (Balasubramanian et al., 1998). This terminal kinase carries out the SIN functions by targeting many essential cytokinetic proteins through phosphorylation (Bohnert et al., 2013; Hergovich, 2016; Sparks et al., 1999; Willet et al., 2019). It also inhibits the Morphogenesis Orb6 Related (MOR) pathway that promotes cell growth (Gupta et al., 2013; Ray et al., 2010; Verde et al., 1998).

We recently identified a novel gene *pkd2* required for the fission yeast cytokinesis (Morris et al., 2019). It encodes the homologue of polycystins, a family of conserved TRP (transient receptor potential) channels (Palmer et al., 2005). Their loss of function mutations lead to one of the most common human genetic disorders, Autosomal Dominant Polycystic Kidney Disease (ADPKD) which manifests through the proliferation of numerous liquid-filled cysts in the kidneys (Hughes et al., 1995; Mochizuki et al., 1996). Most polycystins studied so far are calcium-permissive cation channels sensitive to membrane tension (Gonzalez-Perrett et al., 2001; Hanaoka et al., 2000; Liu et al., 2018; Nauli et al., 2003; Su et al., 2018), but their cellular functions are surprisingly diverse. While the fruit fly and worm polycystins are essential for male fertility (Barr and Sternberg, 1999; Watnick et al., 2003), that of the green algae *Chlamydomonas reinhardtii* plays a non-essential structural role in the primary cilia (Huang et al., 2007; Liu et al., 2020; Wood et al., 2012) and the homologue in the social amoebae *Dictyostelium discoideum* contributes to cell locomotion (Lima et al., 2014). Overall, the mechanism underlying these cellular functions of polycystins remains unclear (for recent reviews see (Hardy and Tsiokas, 2020; Ta et al., 2020)).

Similar to the other unicellular organisms, fission yeast possesses just one polycystin homologue Pkd2p (Palmer et al., 2005). This putative TRP channel localizes to the plasma membrane. While it concentrates at the cell tips during interphase growth, it moves to the equatorial plane during cytokinesis (Morris et al., 2019). It is an essential protein. The null mutant dies either at the single-cell stage or after a couple of rounds of cell division. Depletion of Pkd2 in the mutant *pkd2-81KD* leads to unusually fast constriction of the contractile ring and delayed or failed cell separation (Morris et al., 2019). The cell wall structure remains normal in these Pkd2p depleted cells. In addition to their cytokinesis defects, the mutant cells occasionally shrink but quickly recover. However, the molecular processes underlying these defects remain unknown.

Here we investigated the mechanism of Pkd2p function by characterizing a novel temperature-sensitive *pkd2* mutant and examining its genetic interaction with SIN mutants. We found that this mutation *pkd2-B42* led to strong defects in cell morphogenesis. Most unusual was that the mutant cells frequently shrunk rapidly but always recovered in a process typically lasting no more than 20 minutes. It did not lead to cell lysis and was independent of septation as well as the cell cycle progression. This *pkd2* mutation strongly inhibited cell growth by reducing the tip growth rate and blocking the cell volume extension. Their turgor pressure, modeled through measuring the stiffness of the mutant cells by Atomic Force Microscopy (AFM), was 50% lower than the wild type, but their cell wall appeared to be normal. We identified strong epistatic genetic interactions between *pkd2-B42* and most SIN mutants. The *pkd2* mutations prevented the SIN mutant cells from lysis and increased the septation during cytokinesis. Depletion of Pkd2p also increased the localization of SIN proteins to SPB. We conclude that fission yeast Pkd2p promotes cell expansion likely by regulating the turgor pressure and antagonizes the SIN pathway specifically.

## Results

### A novel temperature-sensitive mutant *pkd2-B42*

To investigate the essential functions of *pkd2,* we set out to isolate a loss of function mutant allele through random mutagenesis. The only available mutant *pkd2-81KD* is a hypomorphic depletion mutant that replaces the endogenous promoter with a weaker one, but 30% of the protein remains (Morris et al., 2019). Through a screen, we identified a novel temperature-sensitive mutant, *pkd2-B42*. It contained eight missense mutations, within the sequence coding for the TRP channel domain (Fig. 1A and Table S1). The mutant cells were inviable at 36°C and above (Fig. 1B). To determine whether the mutant can be rescued by osmotic stabilization of its cell wall, we supplemented the media with sorbitol. Neither 0.6M nor 1.2M sorbitol significantly improved the viability of the mutant at 36°C (Fig. 1C). It was hypersensitive to calcium, similar to *pkd2-81KD* (Fig. 1D). The *pkd2-B42* cells grew more slowly in the liquid media at the restrictive temperature, than either the *wild-type* or *pkd2-81KD* (Fig. 1E). We concluded that *pkd2-B42* is a temperature-sensitive loss of function mutant allele.

**Figure 1.**
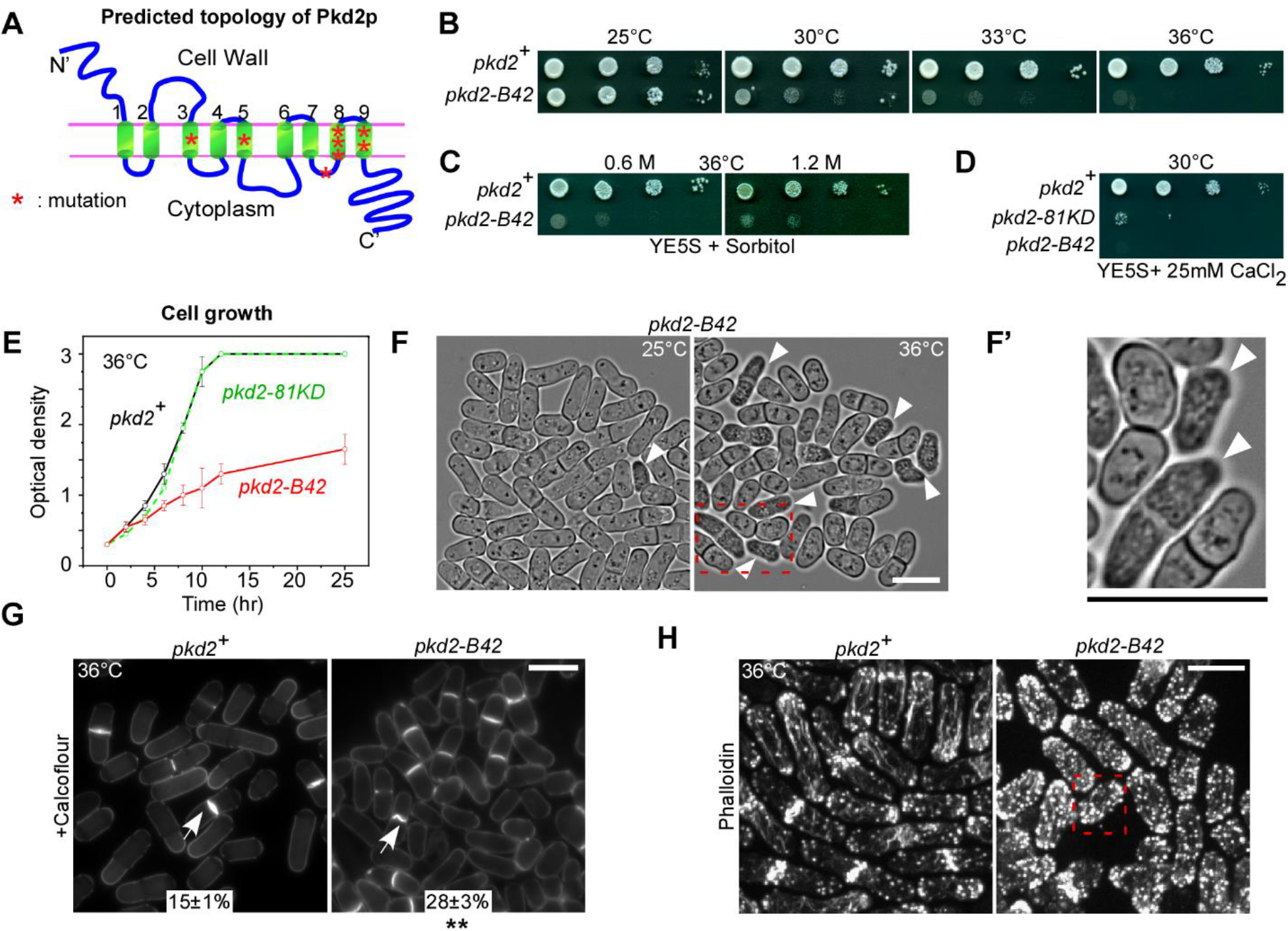
A novel temperature-sensitive mutant *pkd2-B42.* **(A)** Schematic of the predicted topology of the wild-type Pkd2p. The mutated residues in *pkd2-B42* are shown in red asterisks. Numbers denote the nine transmembrane helices. **(B–D)** Ten-fold dilution series of yeast on YE5S plates. **(E)** Growth curve of yeast cultures in YE5S liquid medium at 36°C. Error bars represent standard deviations. **(F)** Bright-field micrographs of *pkd2-B42* cells at either 25°C (left) or 36°C (right). **(F’)** Magnified view of the cells in the rectangle (dashed line). Arrowhead: the shrunk cells. **(G)** Micrographs of wild-type and *pkd2-b42* cells fixed at 36°C and stained with calcofluor to visualize septum. Arrow: septum. Number: the percentage of septated cells (average ± standard deviations, n>500). **(H)** Fluorescent micrograph of fixed and bodipy-phallacidin stained wild-type and *pkd2-B42* cells to visualize the actin cytoskeleton. Scale bars represents 10 μm. Shown are representative data from at least two independent biological repeats. **:P<0.01 (Two-tailed student t-test).

We next examined the morphology of these *pkd2-B42* mutant cells at the restrictive temperature. They were significantly shorter and wider than the wild-type cells (Fig. 1F and Fig. S1A-B). Many (20%) appeared as shrunk and optically opaque, but they were not vacuolated, (Fig. 1F-F’) after being shifted to 36°C for 4 hours. The fraction of such cells continued to increase with extended incubation at the restrictive temperature. Their septation index was one-fold higher than the wild type (Fig. 1G). The septum of some mutant cells appeared as either wavy or slightly curved. Compared to the wild type, the actin cytoskeleton of the mutant appeared disorganized. In particular, the endocytic actin patches were distributed more evenly throughout the cytoplasm, not restricted to either the cell tip or the equatorial plane (Fig. 1H). We concluded that *pkd2-B42* leads to strong defects in cell morphogenesis.

### Deflation of *pkd2-B42* mutant cells

To investigate how the *pkd2* mutant cells became shrunk and opaque, we imaged them at the restrictive temperature with time-lapse microscopy. These mutant cells gained such a unique appearance after rapidly losing their length and width, by 6% (6 ± 2%, average ± standard deviation, n = 38) and 11% (11 ± 4%, n = 26) respectively (Fig. 2A–B, S1C). This transformation usually occurred in less than 5 mins (Fig. 2B). However, such shrinking was temporary. All of them (n =152) gradually regained both their physical dimensions and appearance fully, typically within 15 mins (Fig. 2B). Combined, the whole process lasted ~17 mins (17±7 mins, Fig. 2B). On average, a mutant cell underwent such shrinkage and recovery once every five hours at 37°C (0.19 event per hour per cell, n = 229). We termed this process “Deflation” due to its similarity to the temporary pressure loss of a bike tire.

**Figure 2.**
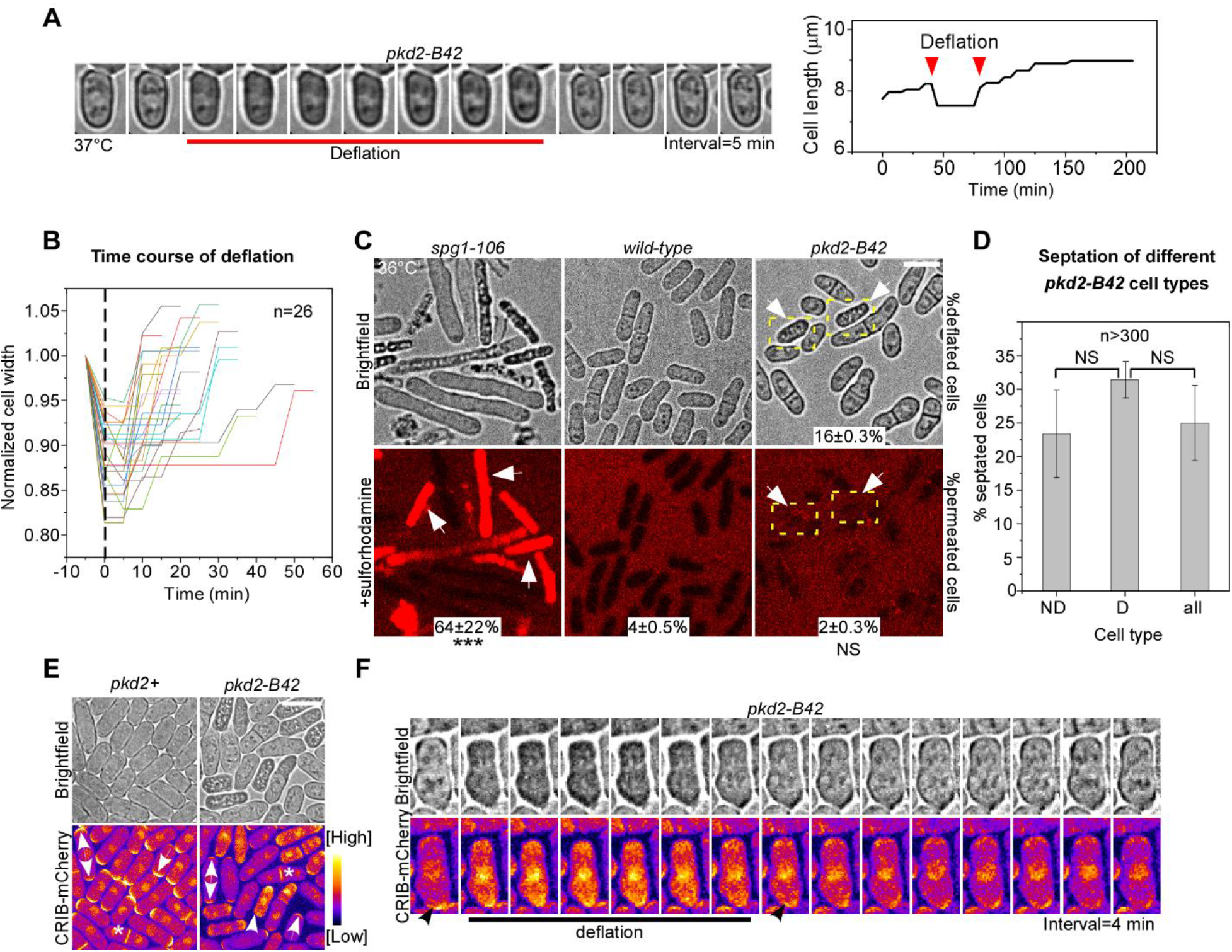
Deflation of *pkd2-B42* cells. **(A)** Deflation of a *pkd2-B42* cell at 37°C. Left: Time-lapse micrographs. Right: Time course of the cell length. **(B)** Time course of normalized width of twenty-six *pkd2-B42* mutant cells during deflation followed by recovery. **(C)** Brightfield (top) and fluorescent (bottom) micrographs of live cells stained with sulforhodamine-B (SRB) to visualize lysed cells respectively in *spg1-106,* wild-type, and *pkd2-B42.* Arrows: deflated *pkd2-B42* cells. Number: percentage of SRB-stained cells (average ± standard deviations, n>400). **(D)** Percentage of septated cells among the non-deflated (ND), deflated (D), and all *pkd2-B42* cells at 36°C, n>300. Error bar represents standard deviation. **(E–F)** Polarity in deflated *pkd2-B42* cells. (E) Brightfield (top) and fluorescent (bottom) micrographs of wild-type and *pkd2-B42* cells expressing CRIB-mCherry at 36°C. Single-headed arrow: monopolar cell. Double-headed arrow: bipolar cell. Asterisk: dividing cell. Arrowhead: deflated cell with mislocalized CRIB-mCherry. **(F)** Time-lapse micrographs of a *pkd2-B42* mutant cell expressing CRIB-mCherry. Interval= 4 min. Representative data from at least two independent biological repeats. Scale bar represents 10 μm. ***: P<0.001, NS: Not significant (Two-tailed student t-test).

Deflation of the *pkd2-B42* mutant cells appeared to be similar to cell lysis of some other fission yeast mutants, prompting us to determine whether they are indeed the same in causing membrane rupture. To test this hypothesis, we stained the cells with sulforhodamine B (SRB), an impermeant dye that fluoresces in the lysed cells (Lemiere et al., 2021; Skehan et al., 1990). This was confirmed through our staining of the *spg1-106* mutant cells (Fig. S1F), many of which (64%) lysed and became fluorescent at the restrictive temperature. In contrast, very few *pkd2* mutant cells (<5%, n >400) were fluorescent in the presence of this dye, similar to the wild type (Fig. 2C). We concluded that deflation of *pkd2-B42* cells does not induce membrane rupture, unlike cell lysis.

The essential role of Pkd2p in cytokinesis (Morris et al., 2019) also prompted us to determine whether such deflation is linked to cell cycle progression including cell separation. First, we determined whether the separation could lead to deflation by quantifying the septation indices of the deflated, non-deflated, and total *pkd2* mutant cells respectively. We found similar percentages of septated cells among the three groups (Fig. 2D). Therefore, deflation is unlikely to result from cell separation. Secondly, we determined whether deflation is linked to the cell cycle progression by measuring the cell length. The length of fission yeast cells increases continuously during interphase before entering mitosis at 14 μm (Mitchison and Nurse, 1985). We found that the length distribution of the deflated *pkd2* mutant cells was similar to that of the whole mutant population (Fig. S1D). Therefore, deflation of *pkd2-B42* cells is cell cycle-independent, not linked to either cell separation or other events in the cell cycle progression.

Lastly, we examined whether deflation is linked to cell polarity. Like most eukaryotes, fission yeast requires the Cdc42-mediated polarity pathway for cell growth and integrity (Miller and Johnson, 1994). We measured the intracellular distribution of active Cdc42p using the fluorescence indicator CRIB-mCherry (Tatebe et al., 2008). Consistent with the previous reports, it localized to either the cell tip or the division plane of the wild-type cells, depending on their cell cycle stage (Fig. 2E). Such polarized localization of CRIB remained unchanged in most *pkd2* mutant cells (Fig. 2E). However, during deflation, the tip and division plane localization of CRIB-mCherry was temporarily lost. Instead, it re-localized to the cytoplasm and nucleus and remained there for ~20 mins (Fig. 2E). Such change followed the initial rapid loss of cell size, but it was gradually reversed as the cells exited deflation (Fig. 2F). We conclude that deflation is correlated with a temporary loss of polarized distribution of active Cdc42.

### Tip and volume expansion of *pkd2-B42* during interphase growth

Morphological defects of *pkd2-B42* cells, as well as their deflation, suggest a likely growth defect. The rod-shaped fission yeast cells grow by extending their tips while maintaining a constant width. We quantified such tip growth through time-lapse microscopy. The mutant cells grew at a rate similar to the wild type at the permissive temperature (Fig. 3A). At the restrictive temperature of 37°C, the wild-type cells maintained their continuous and linear tip extension during interphase growth. In comparison, most mutant cells (66%, n = 229) paused their expansion at least once in a 4-hour window for an average of 17 mins, concurrent with deflation (Fig. 3B, Movie S3). Some (6%) paused 2-3 times. Even between these pauses, the *pkd2-B42* mutant cells extended at a rate of 0.6 ± 0.4 μm hr^-1^ (n = 20), far lower than that of the wild-type cells (~4 μm hr^-1^). Overall, the mutant cells grew at a rate of less than 0.8 μm hr^-1^ (Fig. 3A–B, Movie S1, and S2). Consistent with the relatively normal growth of the depletion mutant *pkd2-81KD* (Fig. 1E), its tip extension rate was similar to that of the wild type (Fig. 3A). We concluded that Pkd2p is required for continuous tip extension during cell growth.

**Figure 3.**
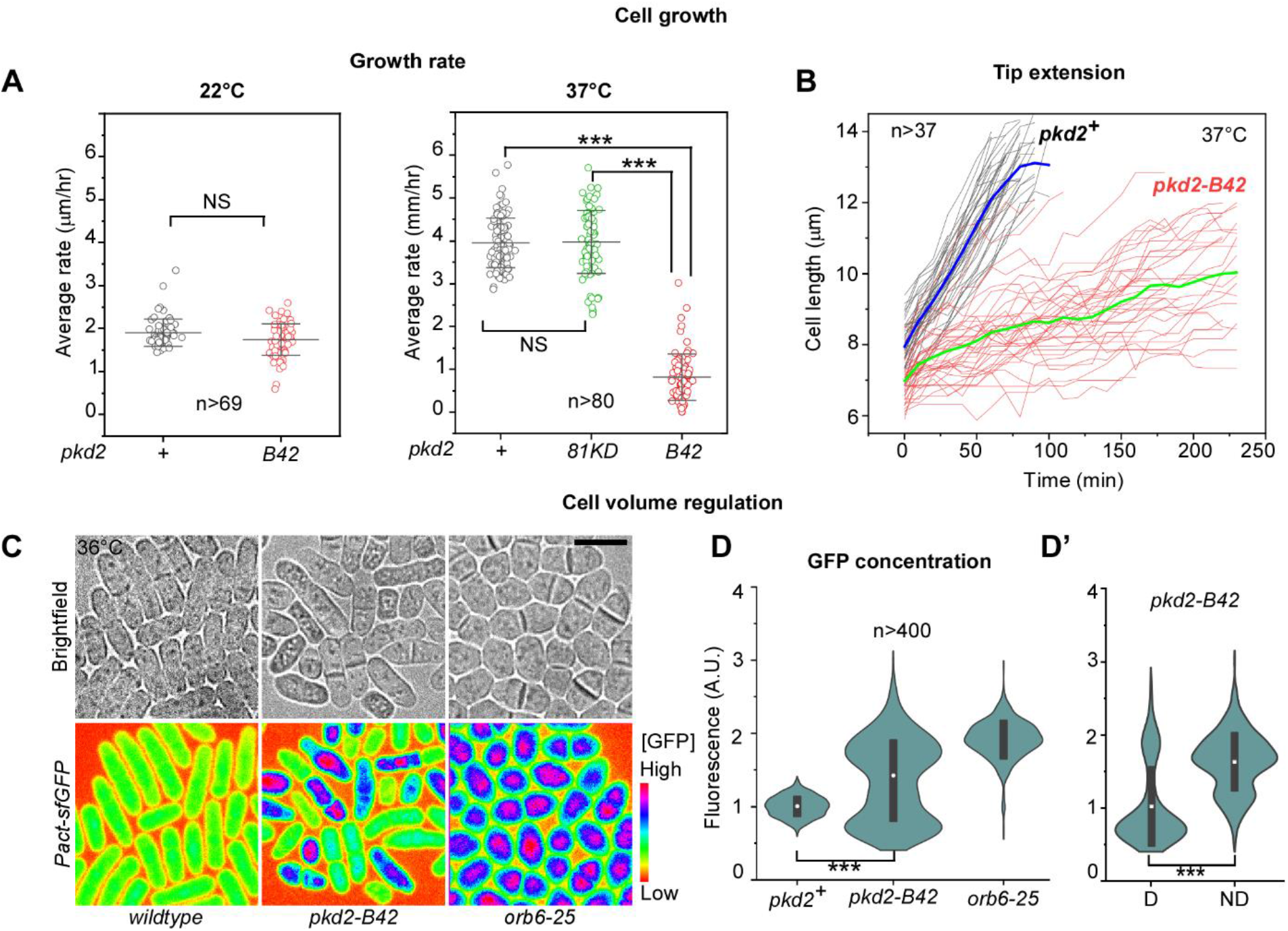
Pkd2p is required for cell size expansion during interphase growth. **(A)** Dot plots of the average tip growth rate of the wild-type *(pkd2^+^)* and the *pkd2* mutant cells at either 22°C or 37°C. Line and error bar represents average ± standard deviation. **(B)** Time courses of the length extension of wild-type and *pkd2-B42* cells at 37°C. Thick lines (blue: wild type, green: *pkd2-B42)* represent the average time courses, n>37. **(C)** Fluorescence micrographs (spectrum-colored) of the wild-type (left),*pkd2-B42* (middle) and *orb6-25* (right) cells at 36°C. All of them constitutively expressed sfGFP. The fluorescence intensity is inversely correlated with the volume expansion rate. **(D)** Violin plots of intracellular GFP concentration. **(D’)** The intracellular GFP concentration of the deflated and non-deflated *pkd2-B42* cells, n>400. The data were pooled from at least two independent biological repeats. Scale bar represents 10 μm. ***: P<0.001, NS: Not significant (Two-tailed student t-test).

The slow growth of *pkd2-B42* cells naturally led to the prediction that they will need a much longer growth period to reach the minimum length required for mitotic entry. The wildtype cells initiate mitosis only after reaching a length of ~14 μm. We tested this hypothesis by staining the cells with DAPI. In an asynchronized population, ~8% of wild-type cells were mitotic. In comparison, we found less than 1% of mitotic cells among the *pkd2* mutant at the restrictive temperature (Fig. S1E). This result confirmed that *pkd2-B42* mutant cells grow very slowly during interphase which delays their mitotic entry.

It remained untested whether the volume of *pkd2-B42* cells expands slowly, considering that the mutant cells are wider than the wild type. Since it is technically challenging to directly measure the volume of a fission yeast cell, we used a novel method that does so indirectly. It quantifies the cellular concentration of a constitutively expressed fluorescence protein which serves as the indirect volume indicator (Knapp et al., 2019). Because the volume of a fission yeast cell expands proportionally with the biosynthesis of such fluorescence proteins like GFP, the intracellular concentration of the fluorescence protein remains constant throughout the cell cycle. In contrast, a defect in the volume expansion would lead to an elevated concentration of the fluorescence protein. Using quantitative fluorescence microscopy, we measured the intracellular concentration of constitutively expressed GFP (super-fold GFP) in both the wildtype and *pkd2-B42* mutant cells. Compared to the wild type, the average concentration of GFP in the mutant cells increased by ~50%, consistent with a defect in the volume expansion (Fig. 3C–D). Similarly, the GFP concentration increased by two-folds (Fig. 3C–D) in another growth-defective mutant *orb6-25* (Verde et al., 1998). Moreover, in contrast to the wild-type cells, the distribution of the cellular GFP concentration was much more heterogenous among the *pkd2* mutant cells (Fig. 3C–D). In particular, the average fluorescence of those in deflation was lower than the others. We conclude that Pkd2p is equally essential for cell volume expansion.

### Turgor pressure of *pkd2-B42* mutant cells

Like the other walled cells, fission yeast cells depend on the turgor pressure, the higher intracellular osmolarity than the environment, to expand. As an indirect measurement of the turgor of the *pkd2-B42* mutant, we quantified its stiffness with atomic force microscopy (AFM) based on a method that we recently established (Gibbs et al., 2021) (Fig. 4A–B). We quantified force-distance curves of surface indentation and cellular spring constants (Fig. 4C). The spring constant of wild-type cells was 42 ± 10 mN m^-1^ (average ± S.D.), consistent with the previously reported values (Gibbs et al, 2021). In comparison, the spring constants of *pkd2-B42* and *pkd2-81KD* mutant cells were 40% lower at the permissive temperature (26 ± 12 mN m^-1^ and 27 ± 9 mN m^-1^ respectively) (Fig. 4D). Interestingly, ‘non-deflated’ *pkd2-B42* cells at 37°C had a cellular spring constant of 28 ± 6 mN m^-1^ (Fig. 4D). In comparison, among ‘deflated’ *pkd2-B42* cells, it was further decreased to 10 ± 5 mN m^-1^ (Fig. 4D). We concluded that the *pkd2* mutant cells were much softer than the wild type.

**Figure 4.**
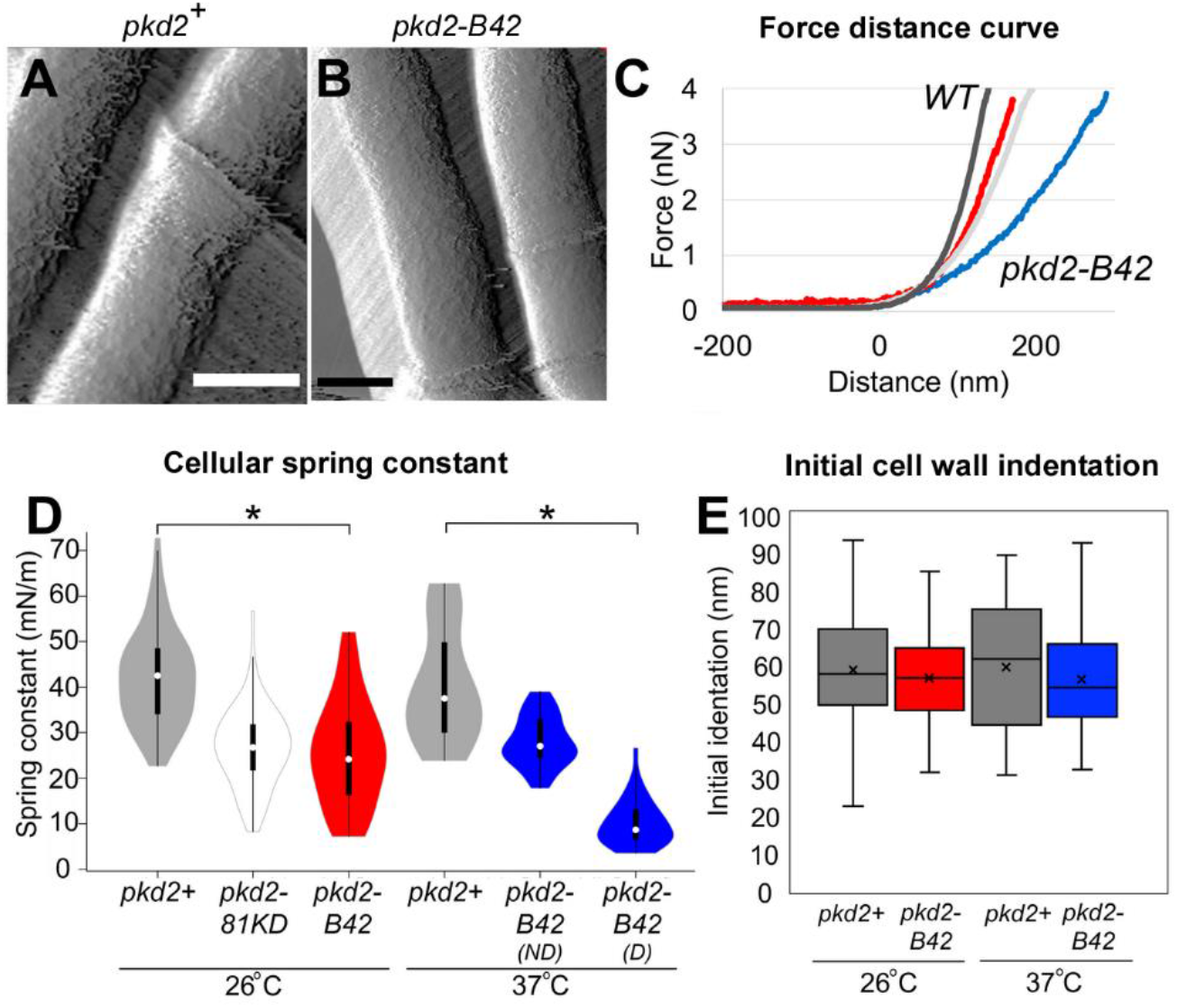
Cellular stiffness is decreased in *pkd2* mutant cells. **(A-B)** Atomic Force Microscope (AFM) deflection images of yeast cells acquired by intermittent contact imaging mode in EMM5S liquid media. **(C)** Representative extension force curves from wild-type (dark grey), *pkd2-81KD* (light grey), *pkd2-B42* at 26°C (red), and *pkd2-B42* at 37°C (blue) cell surface respectively. The horizontal linear region of each curve reflects the cantilever approaching the surface with the zero point along the X-axis indicating the point of contact between the tip and surface. The initial nonlinear deflection of the force-extension curve reflects the initial indentation of the cell wall upon initial cantilever contact. The later linear deflection of the force-extension curve indicates the cellular stiffness. **(D)** Violin plot of cellular spring constants calculated from extension force curves at 26°C and 37°C. ND: not-deflated. D: deflated. White circles show the medians; box limits indicate the 25th and 75th percentiles; whiskers extend 1.5 times the interquartile range from the 25th and 75th percentiles; polygons represent density estimates of data and extend to extreme values. (n > 500 force measurements total from 5 cells per condition). **(E)** Box-and-whisker plot of the initial cell wall indentation from a non-linear region of force-extension curves at 26°C and 37°C. The boxed region indicates the upper and lower quartiles for each data set; the median is indicated by the horizontal line within the box; the mean is indicated by an ‘x’; whiskers extend to high and low data points. (n > 100 force curves evaluated on 5 cells per condition). *: P < 1e^-5^, (ANOVA and student t-tests). Scale bar: 2 μm.

To estimate the decrease in the turgor pressure of *pkd2* mutant cells, we applied the Hertz model (see Equation 2 in Materials and Method) to obtain their Young’s moduli. AFM indentation analysis was used to calculate the Young’s modulus of cellular elasticity and turgor pressure for microbial cells (Arfsten et al., 2010; Gibbs et al., 2021). We modeled the turgor pressure based on the experimentally determined force curves and spring constants. Consistent with previous estimates (Abenza et al., 2015; Atilgan et al., 2015; Davi et al., 2018; Gibbs et al., 2021; Minc et al., 2009), our calculation determined the Young’s modulus of 1.3 ± 0.4 MPa (average ± S.D.) for the wild-type cells (Table S2). The turgor pressure decreased by 50% to 0.6 ± 0.3 MPa in *pkd2-B42* mutant cells, as well as *pkd2-81KD,* at 26°C. The Young’s modulus of deflated temperature-sensitive *pkd2* mutant cells further decreased to 0.3 ± 0.2 MPa at 37°C (Table S2). We concluded that the turgor pressure of *pkd2-B42* is much lower than that of the wild type.

Besides the turgor pressure, another key factor in determining the growth of walled cells is the flexibility of the cell wall (Lew, 2011; Minc et al., 2009). We evaluated this by compressing the yeast cells with the cantilever of AFM. We then calculated the indentation within the nonlinear region of force curves (Arfsten et al., 2010; Arnoldi et al., 2000). At either permissive or restrictive temperature, the indentation values observed for both wild-type and *pkd2-B42* mutant cells were 60–70 nm (Fig. 4E). This was consistent with the thickness of the outermost galactomannan layer of the cell wall measured by electron microscopy (Cortes et al, 2012; Osumi et al, 1998; Osumi et al, 2006). We concluded that the cell wall flexibilities of wild type and *pkd2-B42* are similar.

### Genetic interaction between *pkd2* and SIN mutants

Parallel to the study of the temperature-sensitive *pkd2-B42* mutant, we screened genetic interactions between the available hypomorphic *pkd2-81KD* mutant and a collection of more than thirty cell growth and cytokinesis mutants. We found no genetic interaction with the mutants of either the cell wall synthesis *(bgs4 or ags1)* or the cell wall integrity pathway *(rgf1 or rga7).* The *pkd2* mutant interacted negatively with four cytokinesis mutants, those of *myo2, cdc12, rng2,* and *cdc15.* In contrast, two SIN mutants *sid2-250* and *mob1-R4* stood out for their strong positive genetic interaction with *pkd2-81KD* (Table S2 and (Morris et al., 2019)). Further experiments using seven other SIN mutants (Fig. 5A and Table S2) found that three of them including those of *spg1, sid4,* and *cdc14* interacted similarly with *pkd2-81KD,* particularly at the semi-permissive temperatures of 30°C and 33°C (Table S2, Fig. 5B). Consistent with these positive genetic interactions, we found negative genetic interactions between *pkd2-81KD* and the MOR mutants including *orb6-25* and *mor2-282* (Table S2). MOR is inhibited by SIN during cytokinesis (Gupta et al., 2013; Ray et al., 2010). As expected, in contrast to the above used hypomorphic SIN mutants, the hypermorphic SIN mutant *cdc16-116* (Fankhauser et al., 1993; Furge et al., 1998) interacted negatively with *pkd2-81KD* (Fig. 5C). Like *pkd2-81KD,pkd2-B42* also partially rescued the three SIN mutants that we tested, those of *sid2, mob1,* or *spg1* (Fig. 5D). Our genetic analyses thus suggested an antagonistic relationship between *pkd2* and SIN in cytokinesis.

**Figure 5.**
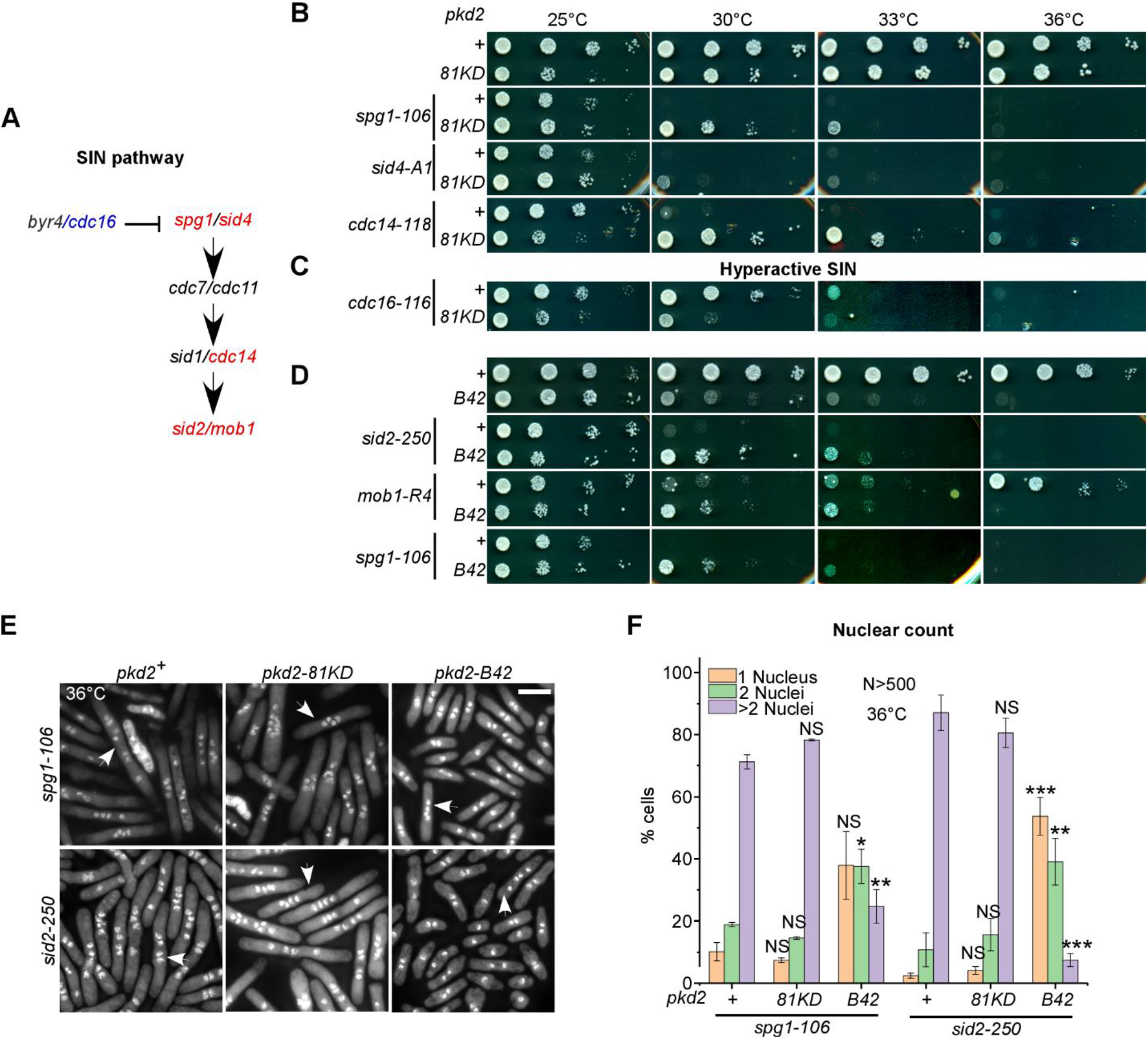
Genetic interactions between *pkd2* and SIN mutants. **(A)** Schematic of the SIN pathway. The genes are colored in either blue (negative genetic interaction with *pkd2-81KD)* or red (positive interaction). **(B–D)** Ten-fold dilution series of yeast on YE5S plates. (B-C) SIN and *pkd2-81KD* SIN mutants. (D) SIN and *pkd2-B42* SIN mutants. **(E–F)** Cytokinesis defects of *pkd2* SIN mutants. (E) Fluorescence micrographs of the mutant cells (36°C) were fixed and stained with DAPI to visualize the nuclei. Arrow: cell with 4 or more nuclei. **(F)** Percentage of mono-, bi-and multi-nucleated cells at 36°C, n > 500. The data were pooled from at least two independent biological repeats. Scale bar represents 10 μm. *:P<0.05, **:P<0.01, ***: P<0.001, NS: Not significant (Two-tailed student t-test).

To understand why the *pkd2* mutations partially rescued the SIN mutants, we characterized the double *pkd2 SIN* mutant cells after a shift to the restrictive temperature for 4 hrs.

First, we counted the fraction of multinucleated cells among the double mutant. The failure of the contractile ring to constrict in the SIN mutants led to a large number (~90%) of bi- or tetra-nucleated cells. The *pkd2-81KD* mutation did not significantly change that (Fig. 5E–F). In comparison, far more SIN *pkd2-B42* mutant cells were bi-nucleate (Fig. 5F), likely undergoing just one round instead of two rounds of cell division in 4 hours. Nevertheless, ~60% of *pkd2-B42* cells were either bi- or tetra-nucleate at 36°C (Fig. 5E–F), far more than the wild type. Interestingly, the fraction of mitotic SIN *pkd2-B42* cells was 5% (Fig. 5E), four-fold higher than that of *pkd2-B42* mutant (1%) and only slightly lower than that of the wild type (8%). Therefore, despite the growth defect of the *pkd2-B42* mutant, most SIN *pkd2* double mutants can enter mitosis but their contractile ring constriction largely fails.

Next, we quantified the septation of the *pkd2* SIN mutants. Most SIN mutants failed to form the septum (Balasubramanian et al., 1998) (Fig. 6A-B, S2A). We found that both *pkd2* mutations partially rescued this defect in the majority of the SIN mutants (Fig. 6A-B, S2A), with the exceptions of the *cdc7* and *sid1* mutants (Fig. S2A). The most significant one was the *cdc11* mutant, in which the septation index increased from 1% to 17% by the *pkd2-81KD* mutation. Nevertheless, many septa in the double mutants appeared to be thin or only partially assembled. We concluded that the *pkd2* mutations significantly promote the septum formation in the SIN mutants.

**Figure 6.**
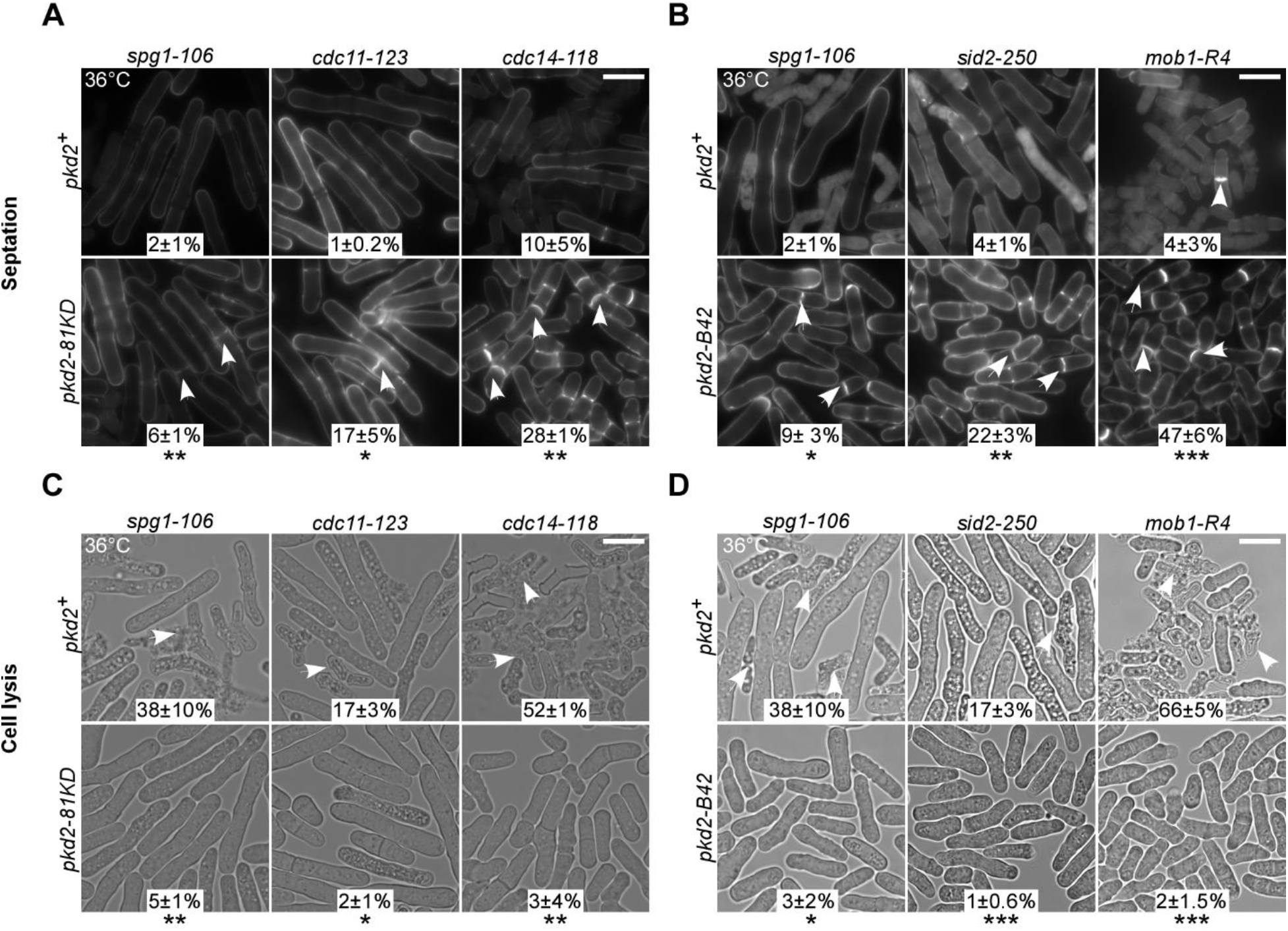
Septation and lysis of *pkd2* SIN mutants. **(A–B)** Micrographs of SIN (top) and SIN *pkd2* (bottom) mutants inoculated at 36°C, fixed and stained with calcofluor. Arrow: septated cell. **(C–D)** Lysis of SIN (top) and SIN *pkd2* (bottom) mutants inoculated at 36°C. Arrow: lysed cell. Number: percentage of either septated **(A-B)** or lysed cells **(C–D) (**average ± standard deviation). The data were pooled from at least two independent biological repeats (n > 400). Scale bar represents 10 μm. *:P<0.05, **:P<0.01, ***: P<0.001, NS: Not significant (Two-tailed student t-test).

Lastly, we measured the cell lysis among the SIN *pkd2* mutants. Although less understood, SIN also ensures the integrity of separating cells and prevents them from lysing (Garcia-Cortes and McCollum, 2009; Gupta et al., 2014; Jin et al., 2006). We confirmed that all SIN mutants lysed in varying degrees ranging from 12% *(sid1-125)* to 52% *(cdc14-118)* (Fig. 6C–D, S2B). The *pkd2* mutants reduced the fraction of lysed SIN mutant cells significantly to less than 5% (Fig. 6C–D, S2B), with the *sid1* mutant being the only exception (Fig. S2B). To determine how specific such rescue was, we tested whether the *pkd2* mutations can prevent other fission yeast mutants from lysis. We found that the *pkd2-81KD* mutant failed to reduce the lysis of *rga7Δ* (Fig. S2B), another cytokinesis mutant that lyses frequently (Liu et al., 2016). Thus, we concluded that the *pkd2* mutations specifically prevent the SIN mutant cells from lysis during cytokinesis.

### Activities and Localization of SIN in *pkd2* mutant

Reduced lysis and increased septation in the *pkd2* SIN mutant cells suggested that the mutations of *pkd2* may have increased the SIN activity. To test this hypothesis, we measured the asymmetric localization of Cdc7p. This SIN pathway kinase distributes asymmetrically between two SPBs during cell division, dependent on the SIN activities (Cerutti and Simanis, 1999; Dey and Pollard, 2018; Feoktistova et al., 2012; Garcia-Cortes and McCollum, 2009; Singh et al., 2011; Sohrmann et al., 1998; Wachowicz et al., 2015). We measured the dwell time of Cdc7p-GFP on the two daughter SPBs (old and new) respectively with time-lapse microscopy. In wild-type cells, Cdc7p-GFP remained at one SPB (new) for ~50 mins, through the end of telophase (Fig. S2B). In comparison, it dwelled on the other SPB (old) for just ~16 mins until the end of anaphase A (Fig. S2B). Both numbers are consistent with the previous measurements (Dey and Pollard, 2018; Garcia-Cortes and McCollum, 2009; Wachowicz et al., 2015). The ratio of the dwell times of Cdc7p on the two SPBs (old:new) averaged ~0.3 for the wild-type cells (Fig. S2C). In comparison, this asymmetry index increased significantly in the *pkd2-81KD* mutant cells to ~0.4 (Fig. S2C), despite that the duration of mitosis did not change significantly (28±2 min vs. 30±4 mins, wild type vs. *pkd2-81KD,* n >65). We concluded that depletion of Pkd2p increases the SIN activity modestly.

In addition, we tested another hypothesis that the mutations of *pkd2* may have enhanced the cytokinetic localization of SIN proteins. We measured the molecular number of Sid2p kinase and its activator Mob1p through quantitative fluorescence microscopy. Both localize to the spindle pole bodies (SPB) in mitosis followed by their translocation to the medial division plane during cytokinesis, before dissipating into the cytoplasm (Salimova et al., 2000; Sparks et al., 1999). We quantified their localization through three steps of cytokinesis, contractile ring assembly and maturation, ring constriction, and cell separation (Fig. 7A) (Wu et al., 2003). In the wild type, each lasts ~30 mins at room temperature, relative to the start of prometaphase marked by the separation of spindle pole bodies (SPBs) (Chen and Pollard, 2011; Morris et al., 2019; Wu et al., 2006) (Fig. 7A). During the first two steps of cytokinesis, the *pkd2-81KD* mutant did not alter the number of these two SIN molecules substantially at either the SPB or the equatorial plane, compared to the wild type (Fig. 7C and S3B). However, during the last step of cytokinesis, the numbers of both SIN molecules doubled at the SPBs of the *pkd2* mutant cells following their exit from the equatorial plane (Fig. 7C–D and S6B–C). We concluded that depletion of Pkd2p significantly increases the localization of SIN proteins at the SPBs.

**Figure 7.**
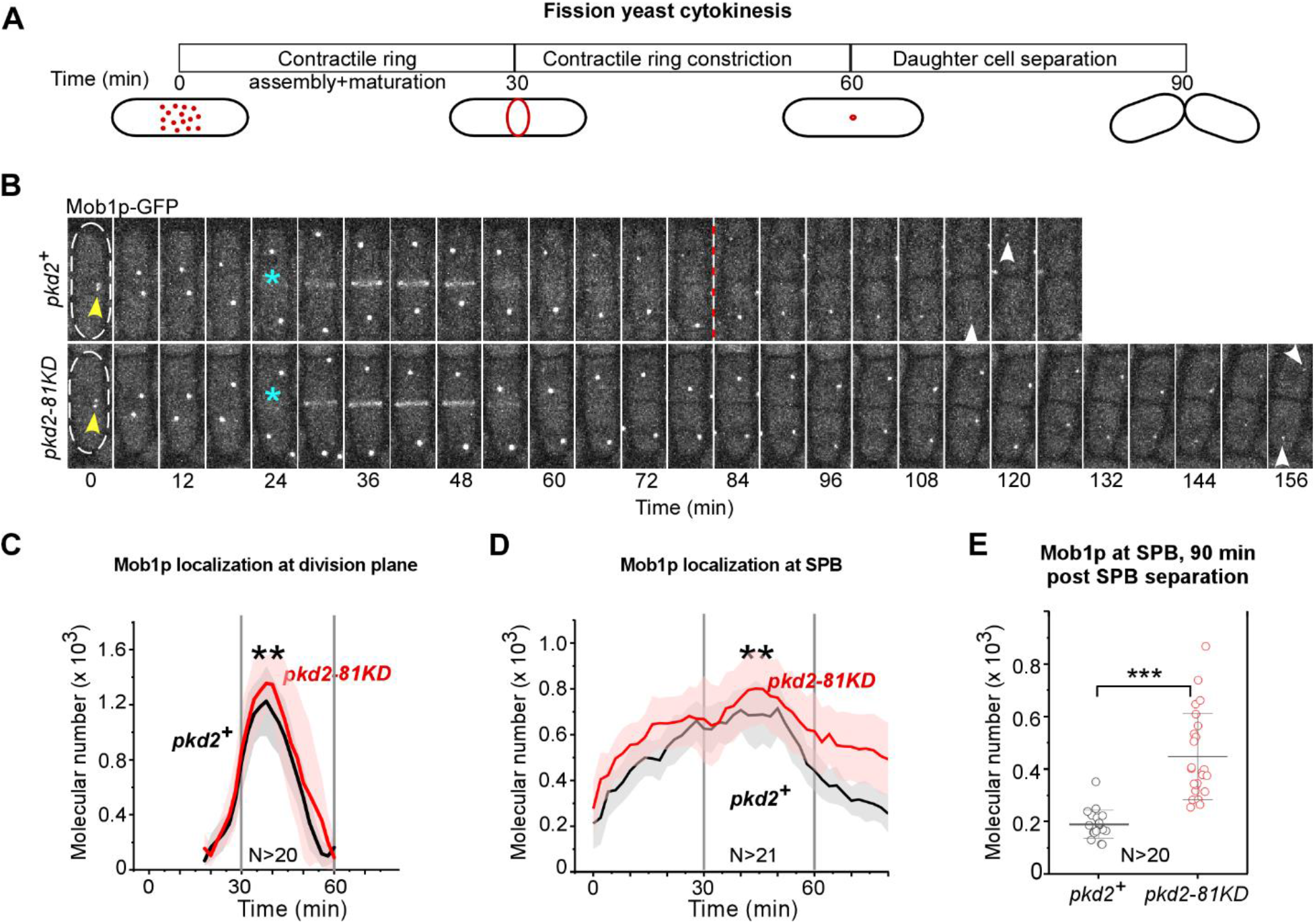
Localization of SIN proteins in *pkd2-81KD* cells. **(A)** Schematic of the average time course of fission yeast cytokinesis. Time zero denotes the separation of SPBs during prometaphase. **(B)** Time-lapse micrographs of either wild-type (top) or *pkd2-81KD* (bottom) cell expressing Mob1p-GFP. Number denotes time in min after SPB separation. Interval = 6 min. Arrowheads: SPB. Asterisk: Appearance of Mob1p-GFP at the division plane. **(C–D)** Average time-courses of the number of Mob1p-GFP (right) molecules at either the division plane **(C)** or SPBs **(D)** of either wild-type (black line) or the *pkd2-81KD* cells (red line). Cloud represents standard deviation. **(E)** Dot plot of the number of Mob1 molecules at 90 mins after SPB separation. Lines and error bar: average ± standard deviation. The data were pooled from two independent biological repeats. ***: P<0.001 (Two-tailed student t-test).

## Discussion

To understand the function of fission yeast polycystin Pkd2 in cytokinesis, we adopted two parallel approaches in this study. First, we isolated a novel *pkd2* loss of function mutant allele. Characterization of *pkd2-B42* revealed surprisingly that Pkd2p is essential for the maintenance and expansion of the cell volume during interphase growth, in addition to its previously reported role in cytokinesis. Secondly, we examined the genetic interplay between *pkd2* and the essential yeast Hippo pathway SIN. This shows that Pkd2p antagonizes both the localization and activity of SIN proteins. Both functions of *pkd2* are likely traced to its activity as a putative cation channel that is sensitive to the membrane tension and permissive to calcium.

### The role of Pkd2 in interphase cell growth

The *pkd2* mutant cells frequently lost and regained their volume in a span of less than 20 mins. Both *pkd2-81KD* and *pkd2-B42* mutants exhibited such a defect of cell growth (Morris et al., 2019), but it was much more prevalent among the latter. This is likely because the depletion mutant *pkd2-81KD* retains ~30% of the endogenous protein. In comparison, the temperature-sensitive mutant cells shall possess very little Pkd2p at the restrictive temperature.

The rapid loss of cell volume during deflation is not due to the plasma membrane rupture. Few *pkd2* mutant cells were stained with the sulforhodamine B dye. It is more likely a result of the outflow of water, similar to the shrinking of cells under hyper-osmotic shock (Poddar et al., 2021). In both cases, the rapid shrinking suggests that the intracellular osmolarity may not have adapted quickly enough to main the turgor pressure. As we have shown in the earlier study, activation of the MAPK kinase Sty1 may have contributed to the graduate recovery following the volume loss (Morris et al., 2019; Toda et al., 1996).

Deflation is independent of cell cycle progression. This also differentiates it from cell lysis that often happens during cell separation in the mutants with a defect in septum biosynthesis (Cortes et al., 2002; Munoz et al., 2013). Interestingly, deflation also occurred during cell division even though there is no cell growth in this period. This suggests that the regulation of turgor is equally important during mitosis and cytokinesis.

To the best of our knowledge, “deflation” or similar phenotype has not been described in other fission yeast mutants. To our surprise, it has not been described in the deletion mutant of *gpd1* which promotes the glycerol biosynthesis for regulating the turgor pressure (Aiba et al., 1995; Minc et al., 2009). Nevertheless, the temporary loss of cell polarity in the deflated *pkd2* mutant cells is reminiscent of such phenotype found in the *gpd1△* cells under hyperosmotic stress (Haupt et al., 2018). Compared to Pkd2p, Gpd1 may be more important for the regulation of turgor under stress. In contrast to the *pkd2* mutants, no such reversible loss of cell volume has been reported in either the mutants of the cell wall integrity pathway (CWI) such as *rgf1△* and *rga7△* (Garcia et al., 2006; Liu et al., 2016) or those of the cell wall sensor *wsc1△* and *mtl2△* (Cruz et al., 2013). These observations suggest that *pkd2* is likely independent from those that sense or repair cell wall damage under stress.

Our data do not support the hypothesis that Pkd2p is required for cell wall biosynthesis. Previously we reported that the cell wall structure of the *pkd2-81KD* mutant cells is similar to that of the wild-type cells (Morris et al., 2019). Moreover, unlike most mutants of cell wall biosynthesis (Liu et al., 1999), the *pkd2* mutant’s viability cannot be fully rescued by supplementing the media with sorbitol. While the mutants of cell wall assembly frequently lysed during cell separation (Cortes et al., 2012; Munoz et al., 2013), the *pkd2* mutant cells rarely did so. We found no genetic interaction between the *pkd2* mutants and the mutants of cell wall synthases either.

Excluding a direct role for Pkd2p in cell wall assembly or repair, the most likely target for Pkd2p is the intracellular osmolarity. In fission yeast as in other fungi, cell expansion depends on the difference between the intracellular and extracellular osmolarity, the turgor pressure (Lew, 2011; Minc et al., 2009). The importance of Pkd2p to turgor pressure is supported by our AFM study showing that the *pkd2* mutant cells are much softer than the wild type. As a putative mechanosensitive channel, Pkd2p may act as a plasma membrane sensor to regulate the intracellular osmolarity and the ensuing turgor pressure.

### The role of Pkd2p in cell division

In light of this newly discovered role of Pkd2p in promoting cell growth, it is surprising that *pkd2* is also needed for cytokinesis because fission yeast cells don’t expand during cell division. One likely explanation is that Pkd2p continues to regulate intracellular osmolarity to maintain cell size even during cytokinesis. This hypothesis is consistent with two cytokinesis defects of the *pkd2* mutant cells (Morris et al., 2019). First, the contractile ring constricts 1.5 times faster in the *pkd2-81KD* mutant. The softer *pkd2* mutant cells may have presented less resistance to the compression force applied by the actomyosin contractile ring during the furrow ingression, compared to the wild type. Second, the *pkd2* mutant cells often failed to separate at the end of cytokinesis. This is also likely due to the mutant’s failure in increasing turgor pressure at the new ends during the cell separation.

### Antagonism between *pkd2* and SIN during cytokinesis

Both *pkd2* mutants partially rescued the SIN mutants by preventing cell lysis and promoting septation, suggesting that Pkd2p plays a direct role in cytokinesis in addition to its function in interphase growth. The lack of interphase growth alone or lack of mitotic entry can’t explain this observation. Unlike the temperature-sensitive mutant *pkd2-B42, pkd2-81KD* has no tip growth defect. Nevertheless, this depletion mutant prevented the SIN mutant from lysis and promoted septum biosynthesis. Further, although *pkd2-B42 SIN* mutant cells exhibited signs of delayed mitotic entry, more than 60% of them still underwent at least one round of cell division without lysis. A likely explanation is that the reduced turgor pressure due to the *pkd2* mutations may indirectly prevent the lysis of the SIN mutant cells.

Unlike the SIN mutants, the mutants of the contractile ring such as those of *myo2, rng2, cdc12*, or *cdc15*, were not rescued by the *pkd2* mutation. In contrast, these cytokinesis mutants exhibited negative genetic interactions with *pkd2-81KD* (Morris et al., 2019). This difference reinforces the idea that the relationship between *pkd2* and SIN during cytokinesis is distinct. It also suggests that reduced turgor pressure due to the *pkd2* mutations alone is not sufficient to rescue the contractile ring constriction.

Reversely, SIN mutations also allow a much larger fraction of *pkd2-B42* mutant cells to progress to mitosis. The mitotic index of *pkd2-B42* SIN mutants is five times higher than *pkd2-B42.* Because fission yeast cells need to reach a minimum length of 14μm to enter mitosis, this observation suggests that the SIN mutations likely promote the tip extension of *pkd2-B42* mutant cells. This also lends support to the proposed antagonistic relationship between SIN and *pkd2.*

Depletion of Pkd2p also increased the SPB localization of the SIN proteins during the cell separation, the last step of cytokinesis. This is despite that Pkd2p localizes at the cleavage furrow throughout cytokinesis (Morris et al., 2019). This putative TRP channel Pkd2p may mediate cytokinetic calcium spikes (Poddar et al., 2021) to activate the calcium signaling pathway in moderating the localization of SIN proteins. Further studies will be needed to determine whether Pkd2p modulates the SIN pathway directly. Alternatively, the antagonism between these two could be indirect through the turgor pressure regulation in cytokinesis.

### Activity and function of polycystin channels

Only mammalian polycystin channels have been characterized for their electrophysiology (DeCaen et al., 2013; Delling et al., 2013; Hanaoka et al., 2000). They are permissive to cations including potassium, sodium, and calcium (Liu et al., 2018), after being activated by mechanical stimuli (Nauli et al., 2003). If the fission yeast Pkd2p channel possesses similar ion-selectivity, it may regulate both ion homeostasis and calcium signaling. For now, the electrophysiology of the Pkd2p channel remains unknown.

Pkd2p homologues can be found among many fungi species, and they constitute one of seventeen essential fungal protein families (Hsiang and Baillie, 2005). It promotes cell growth of *Neurospora crassa* (Bok et al., 2001; Stephenson et al., 2014) through regulating the calcium influx at the cell tip and maintaining a calcium gradient at the expanding hyphae (Silverman-Gavrila and Lew, 2003). The function and mechanism of Pkd2p in cell expansion may be evolutionally conserved in fungi.

In summary, our study discovered that the fission yeast polycystin homologue Pkd2p plays a critical role in the maintenance and expansion of cell size during interphase growth and antagonizes the SIN pathway either directly or indirectly during cytokinesis. We propose that this putative TRP channel regulates both the intracellular osmolarity and the calcium signaling pathway.

## Materials and Methods

### Yeast genetics

Yeast cell culture and genetics were carried out according to the standard methods. YE5S media was used in all experiments unless specified. SPA5S agar plates were used for genetic crosses and sporulation. A SporePlay+ microscope (Singer, England) was used for tetrad dissection and developing different strains. For drop assays, we inoculated the cells in liquid media at 25°C overnight before setting up the ten-fold dilution series on the agar plates. The plates were incubated at designated temperatures for 2-3 days before being scanned by a photo scanner (Epson).

### Identification of the temperature-sensitive *pkd2* mutant

To isolate *pkd2* temperature-sensitive mutants, we employed the marker reconstitution mutagenesis (Tang et al., 2011) with some modifications. First, we amplified the DNA fragment coding for the C-terminal truncated His5 (His5CterΔ) and *ura4* from the pH5C vector (Tang et al., 2011) by PCR using primers P602 and P603 (Table S3). These primers included nucleotide sequences homologous to the 3’UTR of *pkd2.* The amplified DNA fragment was transformed into yeast using the lithium acetate method. The resulting yeast strain (QCY-999) was selected on EMM plates without uracil and confirmed through PCR and Sanger sequencing. Next, the coding sequence of *pkd2* including its 3’UTR was cloned into the vector pH5D (Tang et al., 2011) which contains the missing coding sequence for the C-terminal fragment of His5 (His5Cter). The resultant vector (QCV-193) was used as the template to perform random mutagenesis of *pkd2* through error-prone PCR. The reaction employed Taq DNA polymerase (NEB) in the presence of 200 μM MnCl2 (Sigma). The PCR product consisted of the mutated *pkd2* together with the sequence of His5Cter (PCR product) was transformed into QCY-999 to replace the wild-type gene. The transformants were selected on EMM plates without histidine. The positive clones were further screened for their temperature sensitivity through incubation on the plates supplemented with Phloxin B at 36°C for two days. The temperature-sensitive mutants were isolated and back-crossed with the wild-type twice to confirm their phenotype. The ORF sequence of the *pkd2-B42* mutant was amplified through PCR and sequenced by the Sangers method to identify the point mutations.

### Microscopy

For all microscopy experiments, the cells were inoculated first in liquid media at 25°C for two days before being harvested during the exponential growth phase at a density between 5 x 10^6^ ml^-1^ and 1.0 x 10^7^ ml^-1^. For temperature-sensitive mutants, exponentially growing cell cultures were shifted to the restrictive temperature for 4 hours (unless mentioned) before imaging or fixation.

For microscopy of live cells, the cells were harvested by centrifugation at 4,000 rpm for 1 min. Concentrated re-suspended cells were deposited to a gelatin pad (25% gelatin in YE5S) on a slide. The samples were sealed under the coverslip with VALAP (a mix of an equal amount of Vaseline, lanolin, and paraffin) and imaged.

We used a spinning disk confocal microscope equipped with an EM-CCD camera for fluorescence microscopy. The Olympus IX71 microscope was equipped with the objective lenses 100x (NA = 1.40, oil), 60x (NA = 1.40, oil) and 40x (NA = 1.30, oil), a confocal spinning disk unit (CSU-X1, Yokogawa, Japan), a motorized XY stage and a Piezo Z Top plate (ASI, USA). Solid-state lasers of 405 nm, 488 nm, or 556 nm were used at a power of no more than 5 mW (< 10%). The images were acquired by an Ixon-897 camera controlled by iQ3.0 (Andor, Ireland). We acquired 15 slices at a step size of 0.5 μm or 8 slices at a step size of 1 μm for Z-series. Most time-lapse microscopies were carried out in a dark room whose temperature is maintained at around 23°C. To minimize the temperature variations, we undertook microscopy of both the wild-type and mutants on the same or consecutive days.

To measure the tip growth rate, 20 μL of exponentially growing cell culture was spotted onto a glass coverslip (#1.5) in a 10 mm petri-dish (Cellvis, USA). The coverslip was precoated with 50 μL of 50 μg mL^-1^ lectin (Sigma, L2380) and allowed to dry overnight at 4°C. The cells were overlaid with a 9 mm diameter YE5S agar block (~ 5 mm height) in a covered petri-dish throughout imaging. To measure the growth rate at room temperature, we used the spinning disk confocal microscope. For imaging at 37°C in the temperature-controlled chamber (Precision Control), we used a widE–Field microscope (Olympus IX81) with a 60x oil objective lens (Olympus, NA=1.42) and a CoolSNAP HQ2 camera (Photometrics). Prior to the time-lapse imaging, we shifted the exponentially growing cells to 36°C for inoculation in a water bath for 4 hours.

To measure the septation or mitotic indices, calcofluor or DAPI stained cells were imaged with an Olympus IX81 microscope equipped with a 60x oil lens (NA=1.42) and a digital camera C11440 (Hamamatsu, Japan). Images were acquired using CellSens (Olympus). Brightfield images were illuminated with a LED lamp. For visualizing the cell wall or septum, cells were fixed with 4% paraformaldehyde (Fisher), washed with TEMK buffer, and stained cells with 1μg mL^-1^ of Calcofluor White stain (Sigma). For nuclear staining, cells were fixed with cold 70% ethanol and stained with 1μg mL^-1^ DAPI (Sigma). For some septation indices measurement, we used the spinning disk confocal microscope combined with a 40x oil objective and 405 nm laser.

To visualize the actin cytoskeleton, cells were fixed with 4% paraformaldehyde (Fisher), washed with TEMK buffer, permeated with 1% Triton-X100, and stained with bodipy-phallacidin (Invitrogen).

For staining with Sulforhodamine-B (MP Biomedicals LLC # 152479), 1ml of exponentially growing cells was harvested and resuspended with 1ml of YE5S supplemented with 6.7μg/ml of the fluorescence dye. The cells were immediately spun down and resuspended in ~40μl of supernatant. We dropped 6μl of them onto a glass slide directly and covered them with a coverslip. They were imaged immediately, using the spinning disk confocal microscope with a 40x oil objective and 556 nm laser.

### Image analysis

We used Image J (NIH) to process all the microscopy images, with either freely available or customized macros/plug-ins. For quantitative analysis, the fluorescence micrographs were corrected for both X-Y driftings using StackReg (Thevenaz et al., 1998) and for photo-bleaching using EMBLTools (Rietdorf, EMBL Heidelberg). Unless specified, the average fluorescence of all the Z-slices were used for quantitative measurements. All the measurements were corrected with background subtraction.

The numbers of Sid2p-GFP and Mob1p-GFP molecules were quantified on a calibrated spinning disk confocal microscope (Morris et al., 2019; Wu and Pollard, 2005). The SPB localization was measured by counting the total fluorescence in a 0.8 μm by 0.8 μm (8 x 8 pixels) circle. The background fluorescence was the fluorescence intensities in a ring of 1μm (12 x 12 pixel) surrounding the circle. The contractile ring localization of Sid2p-GFP or Mob1p-GFP was quantified by measuring the fluorescence in a 3 or 4 μm by 0.8 μm (30 or 40 x 8 pixels) rectangle centering on the cell division plane. The background fluorescence was the fluorescence intensity in 3–4 μm by 0.2 μm rectangle (30–40 x 2 pixels) adjoined to the equatorial plane.

The asymmetry of Cdc7-GFP on the two SPBs were measured similarly without counting the molecule numbers due to the relatively weak fluorescence signal.

All the figures were constructed with Canvas X (ACD Systems). The plots were created using either Origin (OriginLab) or KaleidaGraph.

### Atomic Force Microscopy imaging and analysis of cells

Yeast strains were cultured in YE5S medium at 25°C in the mid-log phase (OD_595_ < 0.6) for 36 hours prior to imaging. For restrictive temperature experiments, cells were shifted to 36°C for 8 hours. Cells for AFM imaging and analysis were centrifuged at 500 g and washed with EMM5S medium prior to plating on plastic dishes. Plastic dishes were prepared for yeast cell adhesion by plasma cleaning with PDC-32G plasma cleaner (Harrick Plasma, Ithaca, NY) for 3 minutes followed by incubation for 48 hours at 4°C with 0.8 mg mL^-1^ CellTak adhesive (Corning Life Sciences, Glendale, AZ) diluted in 0.1 M NaHCO3. Dishes were washed twice with ddH2O to remove excess CellTak prior to plating cells. Yeast cells were added to the dish and centrifuged at 500 g for 20 s using a swinging bucket rotor. Non-adherent cells were removed by three EMM5S washes and aspiration.

Topographical height and deflection AFM images were obtained in contact mode using an MFP-3D AFM (Asylum Research, Santa Barbara, CA). Silicon Nitride AFM pyramidal tip PNP-TR probes (Nanoworld, Neuchatel, Switzerland) with a nominal spring constant of 0.32 N m^-1^ were used for all experiments. Cells were maintained in EMM5S media throughout AFM imaging and analysis to prevent drying out. 2D images were exported from MFP-3D software and scale bars were added in Adobe Photoshop. Following topographical imaging, force data was obtained from yeast cells at a deflection setpoint of 4 nN to minimize damage to the cell surface. Ten representative points were selected along the length of each cell, and ten force curves were obtained at each location, for a total of 100 force curves per cell. The extension force curve representing the tip approach and contact with the surface was used to calculate the spring constant for the surface. The spring constants (k_cell_) were calculated from the linear region of the extension curve using a two-spring model with the equation:

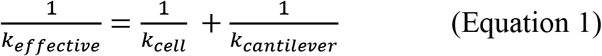

where kcantilever was experimentally calculated using the thermal tuning method (Hutter and Bechhoefer, 1993), and the slope of the extension force curve linear region is keffective (Volle et al., 2008). Data processing was performed through MFP-3D software in Igor Pro to obtain curve fits and spring constants.

AFM extension force indentation curves were used to calculate Young’s elasticity moduli by the Asylum Igor Pro MFP-03D software (WaveMetrics, USA) using the Hertz modeling equation as previously described (Gibbs et al., 2021):

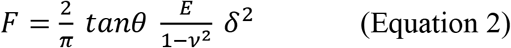

where F is the applied force, θ is the half-opening angle (35°), E is the Young’s modulus, and δ is the indentation depth. A sample Poisson of 0.03 was used in the Hertz modeling.

## Supporting information

Movie S1- wt growth

Movie S2- pkd2-B42 growth

Movie S3- pkd2-B42 deflation

## Abbreviations

(AFM): Atomic force microscopy
(ADPKD): Autosomal dominant polycystic kidney disease
(MOR): Morphogenesis Orb6 related
(SIN): Septation initiation network
(SPBs): Spindle pole bodies

## Acknowledgement

QC conceptualized the study, DS, JG, and QC designed the experiments, DS, DI, EG, MC, JG, and QC carried out the experiments, DS, DI, EG, MC, JG, and QC analyzed the data, DS, JG, and QC wrote the manuscript. The authors thank Abhishek Poddar and Mamata Malla for their technical support. The authors thank Mohan Balasubramanian (University of Warwick, UK), Daniel McCollum (University of Massachusetts Medical School), Jian-Qiu Wu (Ohio State University), and Pilar Perez (University of Salamanca, Spain) for sharing yeast strains and plasmids with us. We acknowledge the National BioResources Project —Yeast Base (Japan) for sending yeast strains. We would like to thank Song-Tao Liu for sharing his microscope.

This work has been supported by the University of Toledo startup fund (QC), National Institutes of Health grant R15GM134496 (QC), DeArce-Koch Memorial Fund (QC), University of Toledo Undergraduate Summer Research and Creative Activities Program (DI), Wellesley College startup fund (JG), and National Science Foundation grant DBI1528288 (JG). The content is solely the responsibility of the authors and does not necessarily represent the official views of the National Institutes of Health. The authors declare no competing interests.

