## Supplemental figures, movie legend and tables for "Fission yeast polycystin Pkd2p promotes the cell expansion and antagonizes the Hippo pathway SIN"

**Supplemental information:**

Figure S1

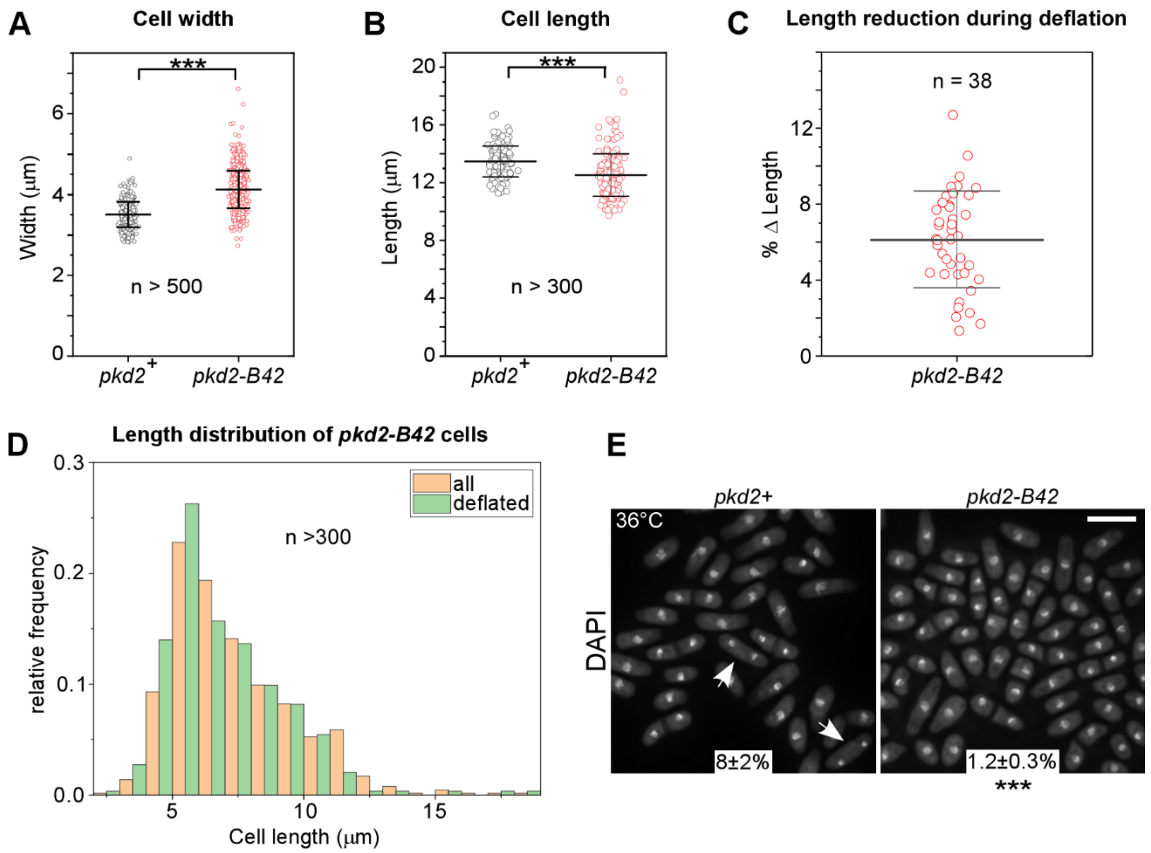


**Supplemental Figure S1. Morphogenesis of the temperature-sensitive mutant *pkd2-B42*. Related to Figure 1 and 2.**

**(A–B)** Dot plots of the width **(A)** and length **(B)** of *pkd2-B42* cells at 36°C, compared to the wild-type (*pkd2+*) n>500 and n>300 respectively. Lines and error bar represents average ± standard deviations **(C)** Dot plot showing reduction in cell length during deflation of *pkd2-B42* cells n=38. **(D)** Histogram showing cell length distribution of *pkd2-B42* cells. Green: deflated *pkd2* mutant cells. Orange: all *pkd2-b42* cells. n>300. **(E)** Fluorescence micrographs of either wild-type (left) and *pkd2-B42* (right) cells at 36°C, fixed and stained with DAPI. Arrow: mitotic cell with condensed nuclei. Number: percentage of mitotic cells (average ± standard deviations, n>500). The data were pooled from at least two independent biological repeats. Scale bar represents 10 µm. ***: P<0.001(Two-tailed student t-test).

Figure S2

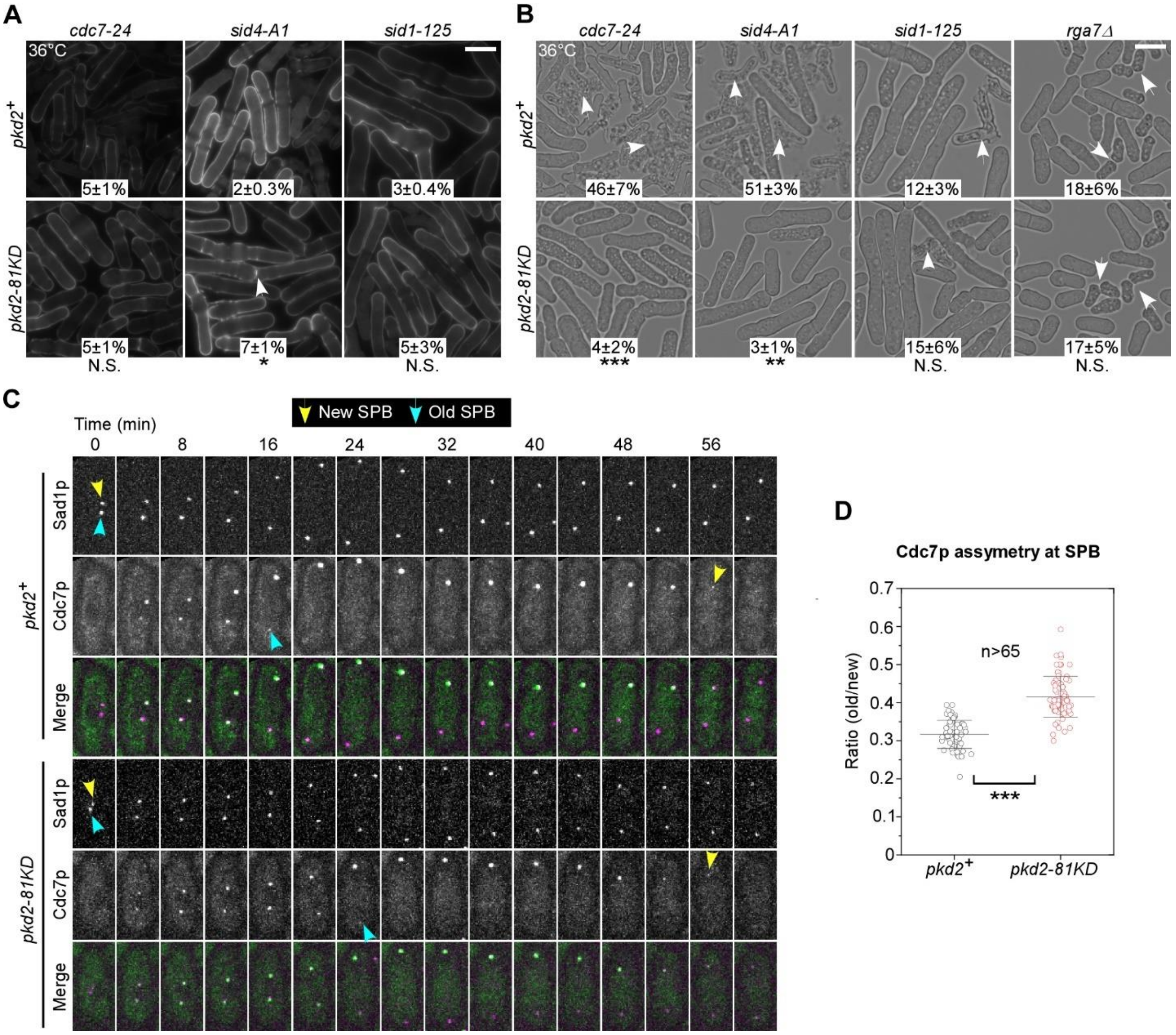


**Supplemental Figure S2. SIN activities in *pkd2* mutant cells. Related to Figure 6**

**(A–B)** Micrographs of fixed SIN (top), SIN *pkd2-81KD* (bottom), *rga7Δ* (top) *rga7Δ pkd2-81KD* (bottom) cells at 36°C. **(A)** Calcofluor stained cells. Arrow: septated cell. **(B)** Arrow: lysed cell. Number: percentages of either septated **(A)** or lysed cells **(B) (**average ± standard deviation, n > 400). (**C–D)** Asymmetric distribution of Cdc7p-GFP at two SPBs during cell division. **(C)** Time-lapse micrographs of either a wild-type or a *pkd2-81KD* expressing both Cdc7p-GFP and SPB marker Sad1-mCherry. Number represents time in mins after the SPB separation. Arrow: new (yellow) or old (blue) SPB. **(D)** Dot plots of ratio (old:new) of the dwelling time of Cdc7p-GFP at the two SPBs, n > 65. Lines and error bar represents average ± standard. The data were pooled from at least two independent biological repeats. Scale bar represents 10 µm. *:P<0.05, **:P<0.01, ***: P<0.001, NS: Not significant (Two-tailed student t-test).

Figure S3

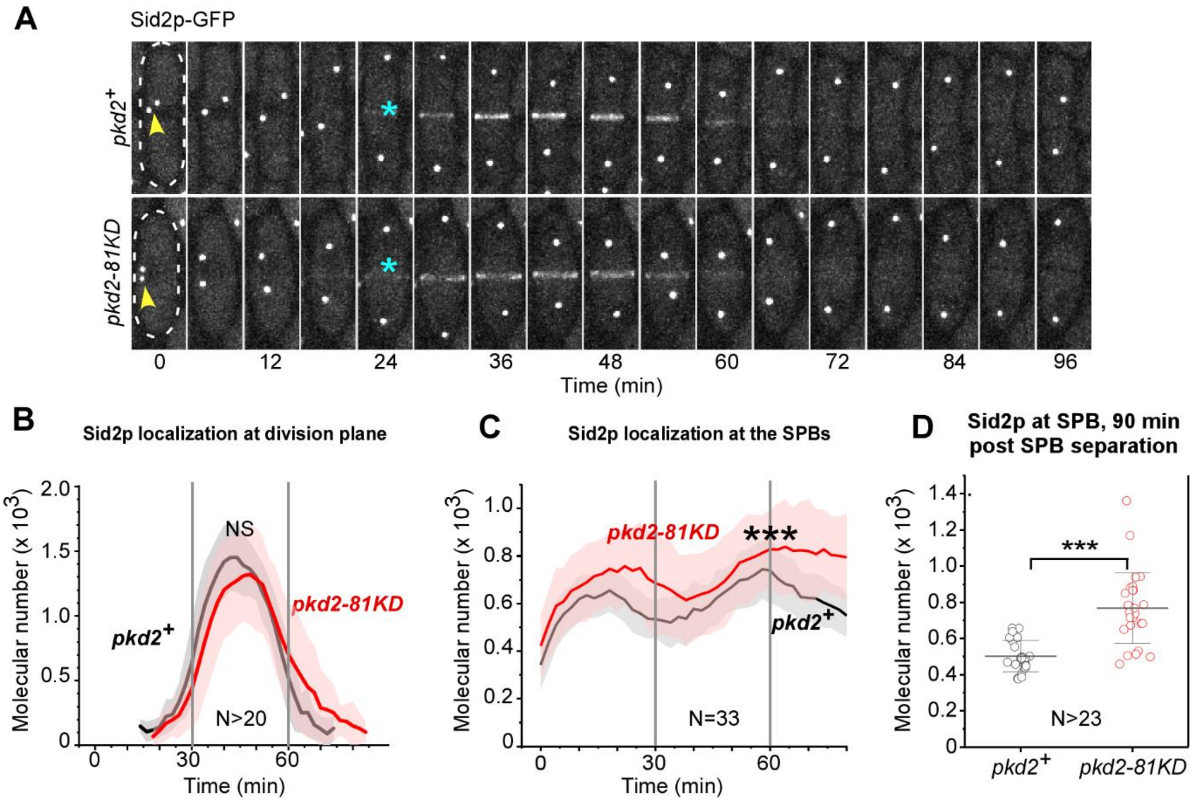


**Supplemental Figure S3. SIN protein localization in *pkd2-81KD* cells during cytokinesis. Related to Figure 7**

**(A)** Time-lapse micrographs of either a wild-type (top) or a *pkd2* mutant (bottom) cell expressing Sid2p-GFP. Number: time in mins after SPB separation. Interval = 6 min. Arrowhead: SPB. Asterisk: appearance of Sid2p-GFP at the division plane. **(B–C)** Average time-courses of the number of Sid2p-GFP molecules in the **(B)** division plane or **(C)** SPBs of either wild-type (black line) or the *pkd2-81KD* cells (red line), n>20 and n=33 for **(B)** and **(C)** respectively. Cloud represents standard deviation. **(D)** Dot plot showing number of Sid2 molecules at 90 min after SPB separation in wild-type or the *pkd2-81KD* cells (red line), n>23. Lines and error bar represents average ± standard The data was pooled from two independent biological repeats. ***: P < 0.001, NS: Not significant (Two-tailed student t-test).

**Supplemental movies:**

**Supplemental movie S1: Time-lapse series of a expanding wild-type cell at 37°C.** Number represents time in minutes after the preceding cell separation**.**

**Supplemental movie S2: Time-lapse series of a expanding *pkd2-B42* mutant cell at 37°C.** Number represents time in minutes after the preceding cell separation.

**Supplemental movie S3: Time-lapse series of *pkd2-B42* cells at 37°C**. Arrow: The deflating cells. Number represents time in minutes.

### Table S1: List of the missense mutations identified in *pkd2-B42*

| Residue | Position in the transmembrane helices | Mutation | Amino acid change |
|---|---|---|---|
| 335 | 3rd | F → S | Hydrophobic to Polar Uncharged |
| 408 | 5$^{th}$ | S → P | Polar Uncharged to Polar Uncharged |
| 521 | Between 7$^{th}$ and 8$^{th}$ | N → D | Polar Uncharged to Negatively Charged |
| 530 | 8$^{th}$ | M → V | Hydrophobic to Hydrophobic |
| 534 | 8$^{th}$ | S → G | Polar Uncharged to Hydrophobic |
| 543 | 8$^{th}$ | Q → E | Polar Uncharged to Negatively Charged |
| 555 | 9$^{th}$ | I → T | Hydrophobic to Polar Uncharged |
| 571 | 9$^{th}$ | I → T | Hydrophobic to Polar Uncharged |

**Table S2: Young's Elasticity Modulus of cellular turgor pressure from Hertz modeling**

| | 26°C | | | 37°C | | |
|---|---|---|---|---|---|---|
| | *pkd2+* | *pkd2-81KD* | *pkd1-B42* | *pkd2+* | *pkd2-B42 (ND)* | *pkd2-B42 (D)* |
| Young's Elasticity Modulus (MPa) | 1.3 ± 0.4 | 0.6 ± 0.3 | 0.6 ± 0.3 | 1.2 ± 0.4 | 0.6 ± 0.3 | 0.3 ± 0.2 |

Note: n > 5 cells evaluated for each condition

Abbreviations: MPa, Megapascal. ND, not deflated. D, deflated.

**Table S3: Genetic interactions between *pkd2-81KD* and other cytokinetic mutants**

| **Gene** | **Mutant** | **Genetic interactions** |
|---|---|---|
| *ace2* | *ace2Δ* | - |
| *agn1* | *agn1Δ* | *No* |
| *ags1* | *mok1-664* | *No* |
| *bgs4* | *cwg1-2* | *No* |
| *cdc11* | *cdc11-123* | *No* |
| *cdc14* | *cdc14-118* | ++ |
| *cdc16* | *cdc16-116* | -- |
| *cdc42* | *cdc42-1625* | *No* |
| *cdc7* | *cdc7-24* | *No* |
| *eng1* | *eng1Δ* | *No* |
| *exo70* | *exo70Δ* | - |
| *mid1* | *mid1Δ* | *No* |
| *mid2* | *mid2Δ* | *No* |
| *mor2* | *mor-282* | -- |
| *myp2* | *myp2Δ* | *No* |
| *orb6* | *orb6-25* | -- |
| *rga7* | *rga7Δ* | *No* |
| *rgf1* | *rgf1Δ* | *No* |
| *rgf2* | *rgf2Δ* | *No* |
| *sid1* | *sid1-125* | *No* |
| *sid4* | *sid4-A1* | + |
| *spg1* | *spg1-106* | ++ |
| *sty1* | *sty1Δ* | *No* |

"++" or "+": strong or weak positive interactions. "--" or "- ": strong or weak negative interactions. No: no interactions. Genetic interactions with the other 11 cytokinesis mutants were published in an earlier study (Morris et al., 2019).

**Table S4: List of the primers used in the screen for *pkd2-B42***

| Number | Sequence |
|---|---|
| QC-P596 | *AGAGTCGAATTTTATTGATG* |
| QC-P597 | *ATGCCGCATAGTTAAGCCAG* |
| QC-P598 | *GTCGTTCTTTTCCTGACATA* |
| QC-P599 | *TATGTCAGGAAAAGAACGAC* |
| QC-P600 | *CTGAAGTCCCAAGCACGAAG* |
| QC-P601 | *CTTCGTGCTTGGGACTTCAG* |
| QC-P602 | *ACGTTTATTAACATTTTATTGAAACAATCTATAGACACCGGTAAGAATAAATCCATAAGCCATTCCCAAA-GGAAATAGTAAGGCTAGTAG* |
| QC-P603 | *GAATATATGCCTATTCGCAATCTAGAATTCCTTTGAATACACCCAATTACAAGCTTAAACGATTCGGTAT-TAAAAATAGGCGTATCACGAGG* |
| QC-P604 | *AGCATATTTGTTGGATGTGC* |
| QC-P605 | *AA-CGATCG-ATGAGGCTTTGGAGAAGCCCAC* |
| QC-P606 | *AA-GGATCC-TTTGGGAATGGCTTATGGATTTATTC* |
| QC-P607 | *GACGAAGCTCTTTCTAGAAGCGTAGT* |

### Table S5: List of fission yeast strains

| Strains number | Genotype | | Source |
|---|---|---|---|
| FY13846 | *h-* | *cps212-282 or mor2-282* | (CP28-2) NBRP Japan |
| FY38515 | *h-* | *his5+:pact1:CRIB[gic2aa1-181]-3mCherry:bsdMX* | (AV2324) NBRP Japan |
| FY527 | *h-* | *leu1-32 ura4-D18 his3-D1 ade6-M216* | Lab stock |
| FY528 | *h+* | *leu1-32 ura4-D18 his3-D1 ade6-M210* | Lab stock |
| JW7551 | *h?* | *cwg1-2 leu1-32* | Jian Wu |
| JW7572 | *h-* | *mok1-664 leu1-32 ura4-D18* | Jian Wu |
| MBY6218 | *h-* | *ade5D ade7::Ade5 his5D leu1-32 ura4D-18* | Mohan Balasubramanian |
| PPG5660 | *h+* | *leu1-32 ura4-D18 cdc42-1425::kanMX6* | Pilar Perez |
| QC-Y569 | *h-* | *cdc11-123 ura4-D18* | Lab stock |
| QC-Y570 | *h-* | *sid4-A1 ura4-D18 leu1-32* | Lab stock |
| QC-Y571 | *h+* | *spg1-106 ura4-D18 leu1-32 ade-210* | Lab stock |
| QC-Y573 | *h+* | *cdc16-116 ura4-D18 leu1-32 ade-M210* | Lab stock |
| QC-Y574 | *h-* | *cdc7-24 ura4-D18 leu1-32 ade-M21X* | Lab stock |
| QC-Y576 | *h+* | *sid2-250 ura4-D18 ade-M21X leu1-32* | Lab stock |
| QC-Y577 | *h-* | *mob1-R4 leu1+ ura4-D18* | Lab stock |
| QC-Y624 | *h+* | *cdc14-118 ura4-D18 ade-216* | Lab stock |
| QC-Y670 | *h-* | *myp2::kanMX6 leu1-32 ura4-D18* | Lab stock |
| QC-Y675 | *h-* | *mob1-mEGFP-kanMX6* | Lab stock |
| QC-Y676 | *h-* | *sid2-mEGFP-kanMX6* | Lab stock |
| QC-Y723 | *h+* | *sid1-125 ura4-D18 leu1-32 ade6-210* | Lab stock |
| QC-Y725 | *h+* | *ace2::kanR ura4-D18 ade6-216 leu1-32 h+* | Lab stock |
| QC-Y743 | *h+* | *far8::kanMX6 ade-M216 ura4-D18 leu1-32* | Bioneer |
| QC-Y803 | *h-* | *orb6-25 leu1-32 ade-M21X* | Lab stock |
| QC-Y810 | *h-* | *KanMX6-P81nmt1-pkd2 leu1-32 ura4-D18 his3-D1 ade6-M21X* | Lab stock |
| QC-Y817 | *h+* | *kanMX6-81xnmt1-pkd2 leu1-32 ura4-D18 his3-D1 ade6-M21X* | Lab stock |
| QC-Y825 | *h-* | *kanMX6-81xnmt1-Pkd2 sid2-250 ura4-D18 ade-M21X leu1-32* | Lab stock |
| QC-Y874 | *h-* | *sty1::kanMX6* | Lab stock |
| QC-Y930 | *h?* | *kanMX6-81xnmt1-Pkd2 sid2-mEGFP-kanMX6* | This study |

| | | | |
|---|---|---|---|
| QC-Y933 | *h?* | *kanMX6-81xnmt1-Pkd2 mob1-mEGFP-kanMX6* | This study |
| QC-Y976 | *h?* | *kanMX6-81xnmt1-pkd2 cdc16-116 ura4-D18 leu1-32 his3-D1 ade-M210* | This study |
| QC-Y977 | *h?* | *kanMX6-81xnmt1-pkd2 cdc14-118 ura4-D18 leu1-32 his3-D1 ade-216* | This study |
| QC-Y978 | *h?* | *kanMX6-81xnmt1-pkd2 cdc11-123 ura4-D18* | This study |
| QC-Y979 | *h?* | *kanMX6-81xnmt1-pkd2 spg1-106 ura4-D18 leu1-32* | This study |
| QC-Y980 | *h?* | *kanMX6-81xnmt1-pkd2 sid4-A1 ura4-D18 leu1-32* | This study |
| QC-Y983 | *h?* | *kanMX6-81xnmt1-pkd2 cdc7-24 ura4-D18 leu1-32 ade-M21X* | This study |
| QC-Y999 | *h-* | *pkd2-3'UTR-his5CterD-ura4* | This study |
| QC-Y1010 | *h+* | *mid1::kanMX6 ade-M216 ura4-D18 leu1-32* | Bioneer |
| QC-Y1011 | *h+* | *mid2::kanMX6 ade-M216 ura4-D18 leu1-32* | Bioneer |
| QC-Y1012 | *h+* | *rga7::kanMX6 ade-M216 ura4-D18 leu1-32* | Bioneer |
| QC-Y1013 | *h+* | *rgf1::kanMX6 ade-M216 ura4-D18 leu1-32* | Bioneer |
| QC-Y1017 | *h?* | *kanMX6-81xnmt1-pkd2 myp2::kanMX6 ade-M21X ura4-D18 leu1-32* | This study |
| QC-Y1022 | *h?* | *kanMX6-81xnmt1-pkd2 mid1::kanMX6 ade-M21X ura4-D18 leu1-32* | This study |
| QC-Y1023 | *h?* | *kanMX6-81xnmt1-pkd2 mid2::kanMX6 ade-M21X ura4-D18 leu1-32* | This study |
| QC-Y1024 | *h?* | *kanMX6-81xnmt1-pkd2 rga7::kanMX6 ade-M21X ura4-D18 leu1-32* | This study |
| QC-Y1025 | *h?* | *kanMX6-81xnmt1-pkd2 rgf1::kanMX6 ade-M21X ura4-D18 leu1-32* | This study |
| QC-Y1031 | *h-* | *pkd2::pkd2-B42-ura4+-his5+ leu1-32* | This study |
| QC-Y1032 | *h+* | *pkd2::pkd2-B42-ura4+-hist5+ leu1-32* | This study |
| QC-Y1032 | *h+* | *pkd2::pkd2-B42-ura4+-hist5+ leu1-32* | This study |
| QC-Y1036 | *h?* | *pkd2::pkd2-B42-ura4+-his5+ leu1-32 ade-M21X sid2-250* | This study |
| QC-Y1037 | *h?* | *pkd2::pkd2-B42-ura4+-his5+ mob1-R4 leu1+* | This study |
| QC-Y1061 | *h+* | *eng1:: KanMx6 ade-M216 ura4-D18 leu1-32* | Bioneer |
| QC-Y1062 | *h+* | *rgf2::kanMX6 ade-M21X ura4-D18 leu1-32* | Bioneer |
| QC-Y1070 | *h-* | *ura4::Pact-sfGFfP-ura4 leu1-32 his3-D1 ade6-M216* | This study |
| QC-Y1072 | *h?* | *pkd2::pkd2-B42-ura4+-his5+ leu1-32 far8::kanMX6* | This study |
| QC-Y1073 | *h?* | *kanMX6-81xnmt1-pkd2 far8::kanMX6 ade-M216 ura4-D18 leu1-32* | This study |
| QC-Y1081 | *h?* | *eng1:: KanMx6 pkd2::pkd2-B42-ura4+-his5+ leu1-32* | This study |
| QC-Y1082 | *h?* | *rgf2:: KanMx6 pkd2::pkd2-B42-ura4+-his5+ leu1-32* | This study |
| QC-Y1085 | *h?* | *eng1:: KanMx6 kanMX6-81xnmt1-pkd2 ade-M21X ura4-D18 leu1-32* | This study |

| | | | |
|---|---|---|---|
| QC-Y1093 | *h?* | *pkd2::pkd2-B42-ura4+-his5+ leu1-32 mok1-664* | This study |
| QC-Y1096 | *h?* | *mok1-664 KanMX6-81xnmt1-pkd2 leu1-32 ura4-D18* | This study |
| QC-Y1108 | *h?* | *cwg1-2 pkd2::pkd2-B42-ura4+-his5+ leu1-32* | This study |
| QC-Y1109 | *h?* | *cwg1-2 leu1-32 kanMX6-81xnmt1-pkd2* | This study |
| QC-Y1162 | *h+* | *agn1::kanMX6 ade-M216 ura4-D18 leu1-32* | Bioneer |
| QC-Y1168 | *h?* | *KanMX6-81xnmt1-pkd2 sid1-125 ura4-D18 leu1-32 ade6-210* | This study |
| QC-Y1178 | *h?* | *KanMX6-81xnmt1-pkd2 ace2::kanR* | This study |
| QC-Y1181 | *h?* | *agn1::kanMX6 pkd2::pkd2-B42-ura4+-his5+ leu1-32* | This study |
| QC-Y1190 | *h?* | *agn1::kanMX6 kanMX6-81xnmt1 pkd2 ura4-D18 leu1-32 ade6-210* | This study |
| QC-Y1198 | *h?* | *orb6-25 pkd2::pkd2-B42-ura4+-his5+ leu1-32 ade-M21X* | This study |
| QC-Y1227 | *h?* | *pkd2::pkd2-B42-ura4+-his5+ leu1-32 his5+:pact1:CRIB[gic2aa1-181]-3mCherry:bsdMX* | This study |
| QC-Y1233 | h? | *ura4::Pact1-pkd2-sfGFP-ura4+ orb6-25 leu1-32 ade-M21X* | This study |
| QC-Y1250 | *h+* | *mor2-282 (cps12-282) pkd2::pkd2-B42-ura4+-his5+ leu1-32* | This study |
| QC-Y1254 | *h?* | *exo70Δ::kanMX6* | This study |
| QC-Y1256 | *h?* | *pkd2::pkd2-B42-ura4+-his5+ leu1-32 exo70Δ::kanMX6* | This study |
| QC-Y1274 | *h?* | *pkd2::pkd2-B42-ura4+-his5+ leu1-32 ade-M21X spg1-106* | This study |
| Qc-Y1307 | *h?* | *cdc42-1625 pkd2::pkd2-B42-ura4+-his5+ leu1-32* | This study |
| QC-Y1317 | h? | *pkd2::pkd2-B42-ura4+-his5+ leu1-32 ura4::Pact-sfGFfP-ura4* | This study |
| QC-Y1352 | *h?* | *sty1::kanMX6 pkd2::pkd2-B42-ura4+-hist5+ leu1-32* | This study |
| QC-Y1408 | h? | *cdc7-GFP::ura4 Sad1-mCherry-NatMX6* | This study |
| QC-Y1409 | h? | *kanMX6-81xnmt1-pkd2 leu1-32 cdc7-GFP::ura4 Sad1-mCherry-NatMX6* | This study |
